# Nitric Oxide Attenuates Human Cytomegalovirus Infection yet Disrupts Neural Cell Differentiation and Tissue Organization

**DOI:** 10.1101/2021.11.09.467865

**Authors:** Rebekah L. Mokry, Benjamin S. O’Brien, Jacob W. Adelman, Suzette Rosas, Allison D. Ebert, Scott S. Terhune

**Author notes:** Corresponding author (SST).

## Abstract

Human cytomegalovirus (HCMV) is a prevalent betaherpesvirus that is asymptomatic in healthy individuals but can cause serious disease in immunocompromised patients. HCMV is also the leading cause of viral-mediated birth defects. Many of these defects manifest within the central nervous system and include microcephaly, sensorineural hearing loss, and cognitive developmental delays. Nitric oxide is a critical effector molecule produced as a component of the innate immune response during infection. Using a 3-dimensional cortical organoid model, we demonstrate that nitric oxide inhibits HCMV spread and simultaneously disrupts neural rosette structures resulting in tissue disorganization. Nitric oxide also attenuates HCMV replication in 2-dimensional cultures of neural progenitor cells (NPCs), a prominent cell type in cortical organoids that differentiate into neurons and glial cells. The multipotency factor SOX2 was decreased during nitric oxide exposure, suggesting early neural differentiation is affected. Maximal mitochondrial respiration was also reduced in both uninfected and infected NPCs. We determined this reduction likely influences neural differentiation as neurons (Tuj1+GFAP-Nestin-) and glial populations (Tuj1-GFAP+Nestin-) were reduced following differentiation. We also observed changes in calcium signaling during exposure to nitric oxide with increased cellular response to ATP (purinergic receptors) and KCl (voltage gated calcium channels). Importantly, nitric oxide could not rescue HCMV-mediated defects in calcium response. Our studies indicate a prominent, immunopathogenic role of nitric oxide in promoting developmental defects within the brain despite its antiviral activity during congenital HCMV infection.

**Author summary:** Human cytomegalovirus (HCMV) infection can result in serious disease to immunocompromised individuals. HCMV is also the leading cause of viral-mediated congenital birth defects. Congenitally-infected infants can have a variety of symptoms, including microcephaly, sensorineural hearing loss, and developmental delays. The use of 3-dimensional (3-D) cortical organoids to model infection of the fetal brain has advanced the current understanding of developmental defects and allowed a broader investigation of the mechanisms behind disease. Here, we investigate the effect of nitric oxide, a critical effector molecule, on cortical development and HCMV infection. We demonstrate that nitric oxide plays an antiviral role during infection yet results in significant disorganization to cortical tissue. Despite inhibiting viral replication in neural progenitor cells, nitric oxide contributes to differentiation defects of these cells and does not rescue functional consequences of HCMV infection on calcium signaling. Our results indicate that immunopathogenic consequences of nitric oxide during congenital infection promote developmental defects that undermine its antiviral activity.

## Introduction

The prevalent betaherpesvirus human cytomegalovirus (HCMV) can cause serious disease during infection. Individuals at higher risk of disease are those that are immunocompromised or receiving immunosuppressive therapies (1). Additionally, congenital HCMV infection occurs in 0.5-1% of all live births and is the leading cause of viral-mediated birth defects as reported for the United States and United Kingdom (2). Congenital infection occurs through primary or secondary infection of a pregnant person and subsequent vertical transmission to the developing fetus. A subset of these infants will exhibit neurological manifestations, either at birth or later in life, that include microcephaly, seizures, sensorineural hearing loss, or other long-term developmental defects (3–5). The mechanisms behind this pathogenesis are not well-defined but involve infection of the central nervous system (CNS) (6–8).

HCMV has wide cell tropism and replicates in various cell types, including fibroblasts, epithelial cells, endothelial cells, trophoblasts, and cells within the CNS such as neural progenitor cells (NPCs) (9). NPCs are primarily found in the subventricular zone of the brain and differentiate into the main cell types of the CNS, including neuron and glial cells (10). NPCs derived from embryonic and induced pluripotent stem cell (iPSC) are susceptible to HCMV infection; however, the stage of differentiation impacts susceptibility with less differentiated cells being more susceptible (11, 12). Infection decreases proliferation and results in abnormal differentiation of NPCs (13–16). Viral protein expression has been demonstrated to contribute to abnormal differentiation both directly and indirectly. HCMV IE1 sequestering of STAT3 to the nucleus results in decreased multipotency factor SOX2 expression (17–19). Liu et al. (14) demonstrated IE1 promotes ubiquitination and proteasomal degradation of the transcriptional regulator Hes1. Calcium (Ca^2+^) signaling, an important process in NPC proliferation and differentiation, is also reduced during HCMV infection in differentiated NPCs (20).

Identifying the mechanisms behind HCMV pathogenesis during fetal development has been largely confined to 2-dimensional cultures due to species specificity until the recent development of iPSC-derived cortical organoids. This 3-dimensional tissue model recapitulates many structural components and the transcriptional profile of the developing fetal brain (21).

This tissue has recently been used to model CNS defects during congenital HCMV infection. We previously demonstrated that cortical organoids infected with HCMV have altered neural layering and disrupted calcium signaling (20). Brown et al. (22) showed that HCMV-infected iPSCs retain the ability to differentiate into cortical organoids but exhibited gross morphological differences compared to mock-infected iPSC-derived organoids. Additionally, expression of genes involved in neurogenesis and brain development are substantially dysregulated during HCMV infection of cortical organoids (23). The use of cortical organoids to model HCMV infection of the developing brain allows more complex investigation into contributing factors during infection such as the innate immune system.

During HCMV infection, innate immune cells produce several nonspecific antiviral molecules, including nitric oxide via the enzyme nitric oxide synthase 2 (NOS2) (24, 25). NOS2 produces picomolar to micromolar levels of nitric oxide (26, 27), resulting in both infected and uninfected cells exposed to nitric oxide at various concentrations. High levels of nitric oxide can inhibit enzymes containing iron-sulfur clusters, including complex I and complex IV of the electron transport chain (ETC) and aconitase, an enzyme in the TCA cycle (28–30). These inhibitions impact several aspects of cellular function such as proliferation, differentiation, metabolism, mitochondrial function, and energy production. In plated NPCs from mice, nitric oxide can enhance or inhibit proliferation in a concentration-dependent manner (31, 32). Further, Bergsland et al. (33) demonstrated that neuron differentiation was decreased following nitric oxide exposure. However, the full impact of nitric oxide on events within the developing brain is largely undefined.

There is evidence that nitric oxide controls viral infections within the brain while also contributing to disease. Nitric oxide inhibits many RNA and DNA viruses (34), including herpesviruses such as herpes simplex virus-1 (HSV-1), herpes simplex virus-2 (HSV-2), and Epstein-Barr virus (EBV) (35–38). Despite this antiviral activity, nitric oxide promotes immunopathogenesis during infection. During HSV-1 infection, exposure to a nitric oxide donor contributed to amyloid beta (Aβ) accumulation in both neuronal cultures and mouse brains (39). In mice with West Nile virus-induced encephalitis, inhibition of nitric oxide production with a selective NOS2 inhibitor increased survival (40). Wang et al. demonstrated that the conditioned media from Zika virus-infected microglia, which contains nitric oxide, decreased neuronal differentiation (41).

Nitric oxide is also inhibitory to HCMV replication and plays an important role in regulating infection (42–46). A case study described a fatal case of HCMV infection in an individual with NOS2-deficiency despite evidence of past pathogenic infections that did not prove fatal (47). We have recently demonstrated that nitric oxide inhibits HCMV replication in human fibroblasts and retinal pigment epithelial cells through a multi-factorial mechanism involving inhibition of mitochondrial respiration and dysregulation of metabolism (48). It is unknown how these mechanisms impact infection within the fetal brain. Congenitally infected fetal brains have necrotic foci with infiltrating macrophages and microglia that are capable of producing nitric oxide (7, 24, 25). In murine models of congenital CMV infection, embryonic mice and pups also have infiltrating NOS2-expressing macrophages within the brain (49, 50).

Nitric oxide is a potent antiviral yet is considered a double-edged sword due to associated tissue damage. Its impact on CNS development during HCMV infection is unknown. Here, we determine the effect of nitric oxide on HCMV infection of cortical organoids and NPCs as well as immunopathogenic consequences on uninfected cells. We demonstrate that nitric oxide attenuates viral spread in cortical organoids and NPCs while dysregulating neural differentiation. Mitochondrial function and intracellular calcium (Ca^2+^) signaling, important regulators of differentiation, were compromised during nitric oxide exposure. Importantly, nitric oxide does not protect HCMV-mediated defects in Ca^2+^ responsiveness, which is necessary for neural development and function. We demonstrate the role of nitric oxide in controlling HCMV infection while causing immunopathogenic effects that are detrimental to the developing fetal brain. These discoveries provide novel insights into neuropathogenesis observed during congenital HCMV infection.

## Results

### Nitric oxide reduces HCMV spread in cortical organoids

Modeling defects in the CNS during congenital HCMV infection have been largely limited to 2-dimensional cell culture systems due to species specificity. However, the emergence of 3-dimensional cortical organoids has allowed investigation of congenital infection in a tissue model with features of fetal forebrain (20, 22, 23). Congenitally HCMV-infected fetuses have necrotic regions in the brain that contain infiltrating macrophages and microglia (7). These immune cells produce nanomolar to micromolar levels of nitric oxide via NOS2 (24–26), suggesting that nitric oxide is present in the fetal brain during congenital infection. However, the impact of nitric oxide on HCMV infection and tissue within the fetal brain is unknown. To determine the role of nitric oxide during HCMV infection in developing neural tissue, we began our studies by defining the effect of HCMV and nitric oxide on cortical organoids. We have previously demonstrated that nitric oxide inhibits HCMV replication in cultured human fibroblasts and epithelial cells using the spontaneous-release nitric oxide donor diethylenetriamine NONOate (DETA/NO) to mimic nitric oxide production by NOS2 (48, 51). Since macrophages and microglia are not typically present in cortical organoids, we used DETA/NO to determine the impact of nitric oxide on this tissue. Cortical organoids were differentiated from an iPSC line derived from a heathy individual (52, 53). Organoids were cultured to day 35 of development at which time they have developed defined brain region identities (54). Organoids were mock-infected or infected using HCMV strain TB40/E encoding enhanced green fluorescent protein (TB40/E-eGFP) at an MOI of 500 infectious units per microgram of tissue (IU/μg). Viral stocks were produced using MRC-5 human fibroblasts (TB40/EFb-eGFP). Organoids were treated at 2 hours post infection (hpi) and every 24 h with 400 μM DETA/NO or vehicle. As an additional control, organoids were treated with spent donor (DETA) to account for effects of nitric oxide oxidation products (i.e., nitrite and nitrate) and the parental backbone molecule. DETA was prepared by incubating medium containing 400 μM DETA/NO at 37°C for 72 h. GFP expression was not observed in mock-infected organoids at 11 days post infection (dpi) (Fig. 1A). We observed an increase in GFP fluorescence and spread from 4 dpi to 11 dpi in vehicle and DETA-treated, HCMV-infected organoids, (Fig. 1A), indicating efficient HCMV replication. In DETA/NO-treated, HCMV-infected organoids, GFP was dramatically decreased compared to vehicle and DETA (Fig. 1A). We quantified the mean fluorescent intensity (MFI) normalized to the cross-sectional surface area and observed an average 79% decrease during DETA/NO exposure compared to vehicle (Fig. 1B). DETA/NO-treated organoids maintained a similar size to vehicle and DETA conditions, suggesting that nitric oxide did not impact organoid growth (Fig. 1B). These data demonstrate that nitric oxide, but not DETA backbone or nitric oxide oxidation products, reduces HCMV spread in cortical organoids.

**Fig 1.**
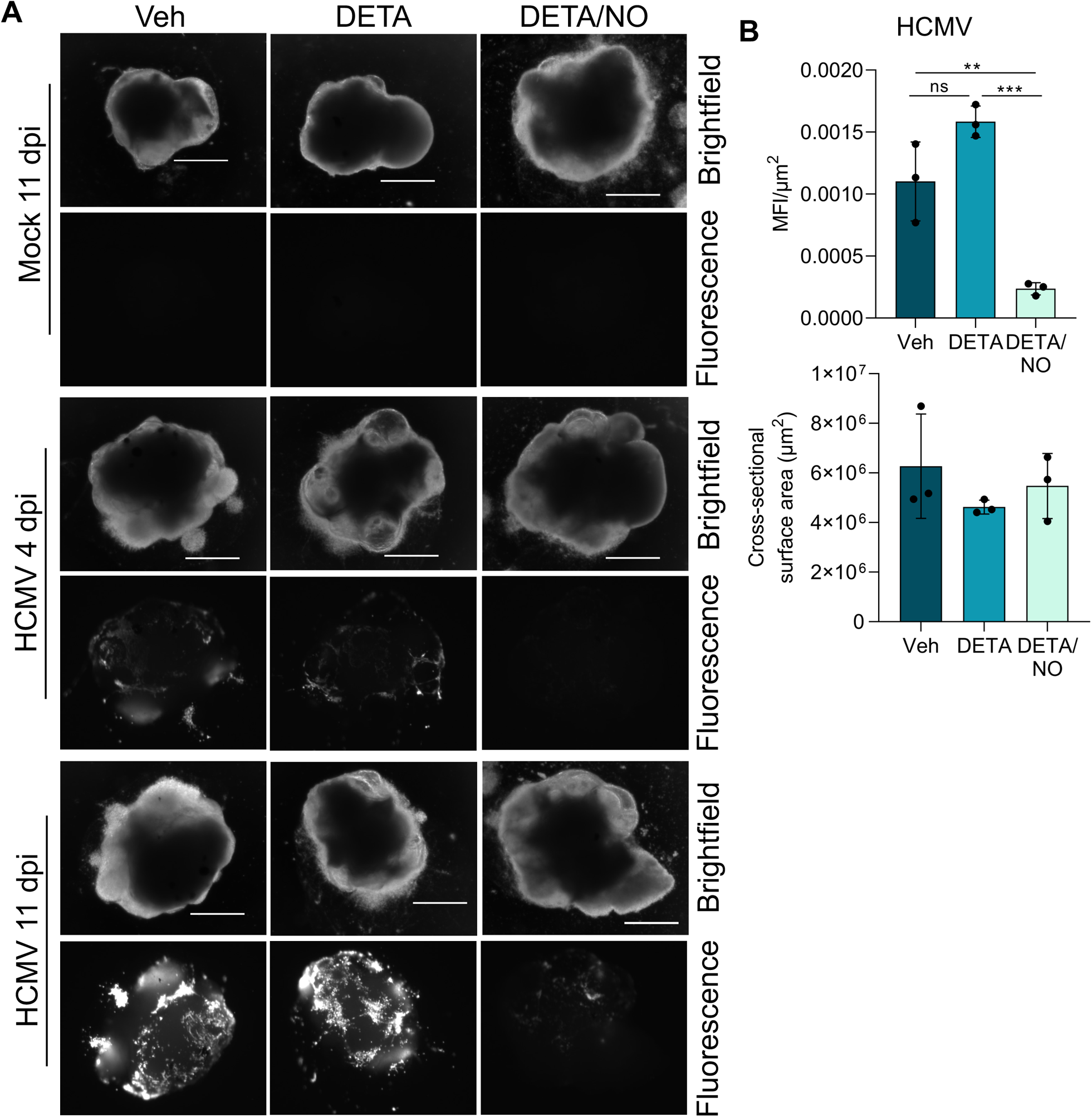
Nitric oxide reduces HCMV spread in cortical organoids. **(A)** Cortical organoids were infected with HCMV strain TB40/E encoding free green fluorescent protein (TB40/E-GFP) propagated in MRC-5 fibroblasts (TB40/EFb-GFP) at an MOI of 500 infectious units/μg (IU/μg) or mock infected, and tissues were treated at 2 hpi with 400 uM of the nitric oxide donor diethylenetriamine NONOate (DETA/NO), spent donor (DETA), or vehicle control. Treatment was changed every 24 h. Images were acquired at 11 dpi using a 1x objective and represent 3 biological replicates per condition with individual organoids considered a biological replicate. Scale bar is 1000 μm. **(B and C)** The cross-sectional surface area (μm^2^) of the organoids and mean fluorescence intensity (MFI) were measured using NIS-Elements software and relative MFI to μm^2^ was quantified. Errors bars represent standard deviation from the mean (SD). One-way ANOVA with multiple comparisons was used to determine significance. *P* = 0.0044, **; *P* = 0.0004, ***; ns = not significant.

### Neural rosette structures and tissue organization are disrupted in nitric oxide-exposed cortical organoids

Similar to the human brain, cortical organoids develop 3-dimensional structure with distinct multicellular layer identities (54). HCMV infection of cortical organoids increases cell death and disrupts tissue morphology and organization, specifically neural rosette structures (20, 22, 23). To determine if nitric oxide inhibition of HCMV spread could alleviate this disruption, we evaluated cell viability and neural rosette structure in cryosectioned organoids using immunofluorescence. Consistent with our previous studies (20), we observed that GFP fluorescence was largely confined to the periphery of the organoid with limited spread to the inner layers (Fig. 2A). We first labeled for fragmented DNA, which is indicative of cell death, using terminal deoxynucleotidyl transferase dUTP nick end labeling (TUNEL) (Fig. 2A). We quantified the TUNEL relative to Hoechst total DNA signal (Fig. 2B). In uninfected organoids, TUNEL/Hoechst ratio was significantly increased by 2.3-fold in DETA/NO-treated organoids compared to vehicle, which did not occur in DETA conditions (Fig. 2B). We observed an increased ratio of TUNEL/Hoechst across all conditions in HCMV-infected organoids (Fig. 2B). These data indicate that nitric oxide increases DNA damage, and possibly cell death, in uninfected organoids but not in HCMV-infected organoids.

**Fig 2.**
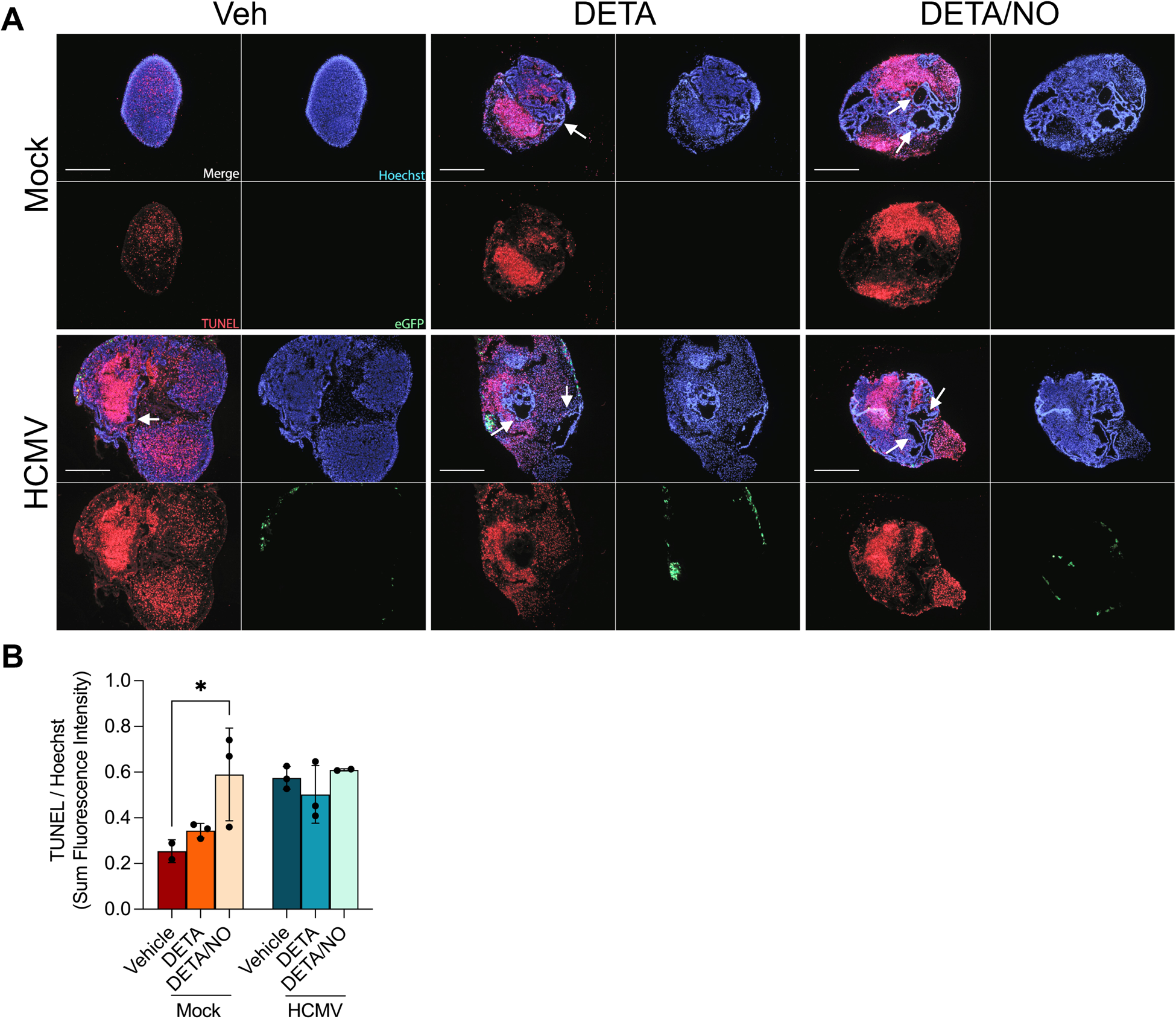
Nitric oxide increases cell death in uninfected organoids but does not further increase cell death in HCMV-infected organoids. Immunofluorescent images of cortical organoid sections at 12 dpi. **(A)** Sections were labeled for terminal deoxynucleotidyl transferase dUTP nick end labeling (TUNEL) (red) with Hoechst (blue) used to stain nuclei. HCMV-infected organoids show GFP expression (green) with no expression observed in mock-infected organoids. Scale bar is 500 μm. Images represent organoid sections from two to three organoids per condition. **(B)** The sum fluorescent intensity ratio of TUNEL to Hoechst (TUNEL/Hoechst) was quantified from two to three organoid sections per condition. Error bars represent SD. Two-way ANOVA with multiple comparisons was used to determine significance. *P* = 0.02, *.

Tissue sections also revealed substantial changes to organoid organization, including regions absent of TUNEL and Hoechst (Fig. 2A). The structures were present in all organoid conditions but were more evident in mock and HCMV-infected organoids exposed to nitric oxide (Fig. 2A, arrows). As the regions appeared similar to radial neural rosette structures formed by SOX2-expressing neural progenitor cells (NPCs), we labeled for SOX2, a transcription factor necessary for progenitor maintenance and a marker for NPCs (55–58). We observed that mock-infected, vehicle-treated organoids displayed SOX2-expressing cells organizing in layered radial neural rosette morphology (**Fig. 3A** and **B**, arrowheads). As previously described (20), rosette structures were disrupted in HCMV-infected, vehicle-treated organoids (**Fig. 3A** and **B**, arrows). SOX2-expressing cells also formed rosette-like structures in nitric oxide exposed cortical organoids. However, the structures lacked the morphology and layering of a typical rosette, and the overall organization of the structures were compromised (Fig. 3A, arrowheads and arrows). DETA alone in mock and HCMV-infected organoids exhibited an intermediate phenotype with some sections displaying more typical rosette morphology and others showing disrupted rosette-like structures (**Fig. 3A** and **B**, arrowheads and arrows). Our data indicate that nitric oxide, and possibly its oxidation products, disrupt cortical organoid organization. Overall, these results suggest that nitric oxide inhibits viral spread in cortical organoids yet disrupts neural rosettes and organization regardless of infection.

**Fig 3.**
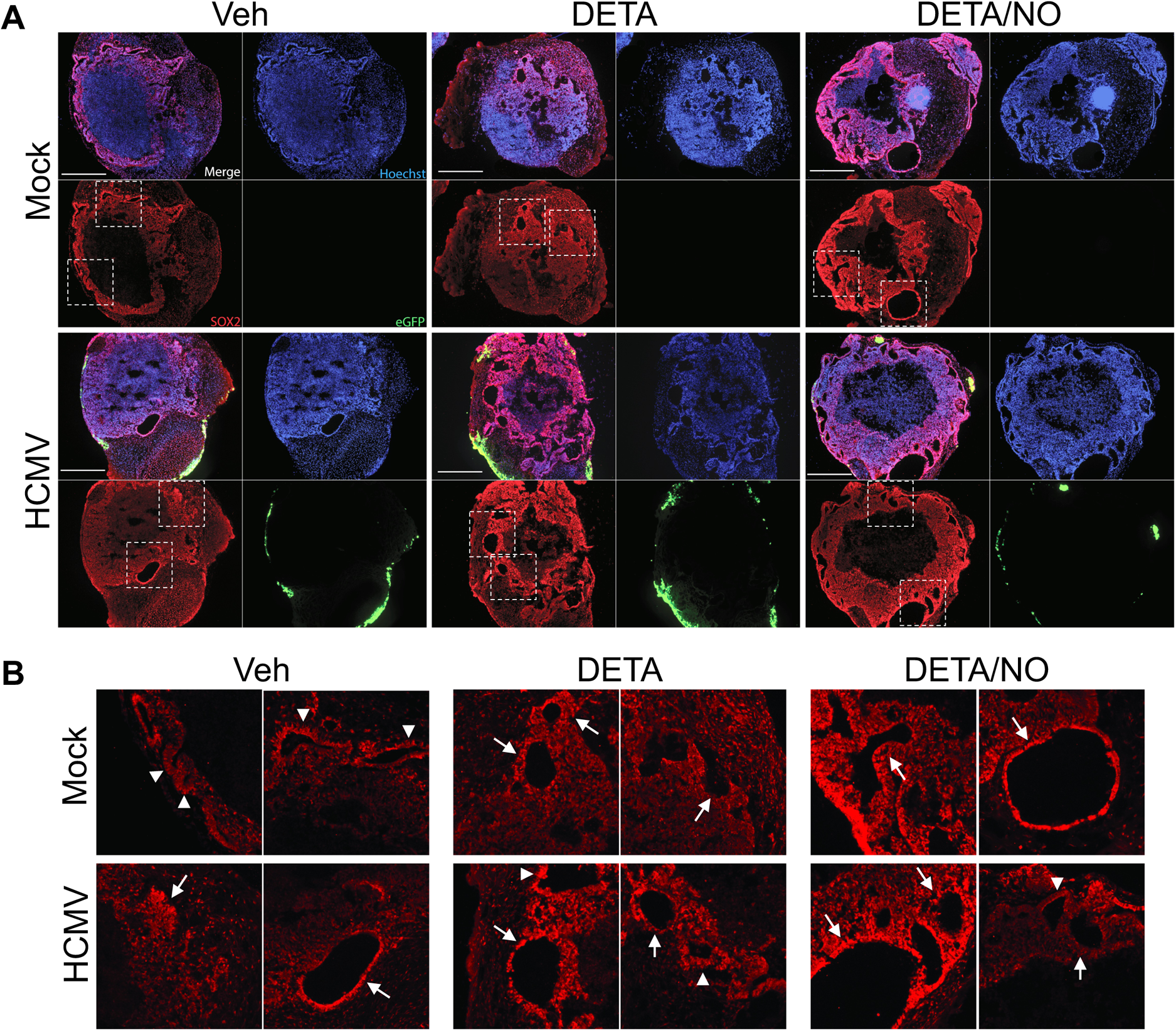
Neural rosette structure and tissue organization are disrupted in cortical organoids during nitric oxide exposure. **(A)** Organoid sections were labeled for SOX2 (red) with Hoechst (blue) used to stain nuclei. White boxes indicate regions enlarged in B. Scale bar is 500 μm. **(B)** Arrowheads point to SOX2-expressing cells organizing in radial neural rosette structures. Arrows point to disorganized rosette-like structures. Images represent organoid sections from three organoids per condition.

### Nitric oxide attenuates HCMV replication and alters markers of NPC proliferation

SOX2-expressing NPC rosettes were disrupted in nitric oxide exposed organoids (Fig. 3A and B). Therefore, we explored the impact of nitric oxide on NPC development and function during infection. NPCs are susceptible to HCMV infection and are significantly disrupted in HCMV-infected organoids (20, 22). We used 2-dimensional cultures of NPCs differentiated from the same healthy iPSC line from which the cortical organoids were derived. NPCs were cultured as neurospheres before dissociation and plating (59). Initially, we examined the impact of nitric oxide on HCMV replication. To increase infection efficiency, we used TB40/E-eGFP produced in ARPE-19 epithelial cells (TB40/EEpi-eGFP) (60). NPCs were plated, incubated for 3 days, and infected with TB40/EEpi-eGFP at an MOI of 3 infectious units per cell IU/cell. At 2 hpi, cultures were treated with 200 μM DETA/NO, DETA (spent donor), or vehicle control that were replaced every 24 h. Viral DNA levels were quantified at 2, 48, and 96 hpi (Fig. 4A). Viral titers from 96 hpi were determined on ARPE-19 epithelial cells (Fig. 4A). Viral DNA levels were reduced by 0.6-log and viral titers by 0.9-log at 96 hpi during nitric oxide exposure. Our data indicate nitric oxide attenuates HCMV replication in NPCs, which is consistent with our previous studies in human fibroblasts and epithelial cells (48).

**Fig 4.**
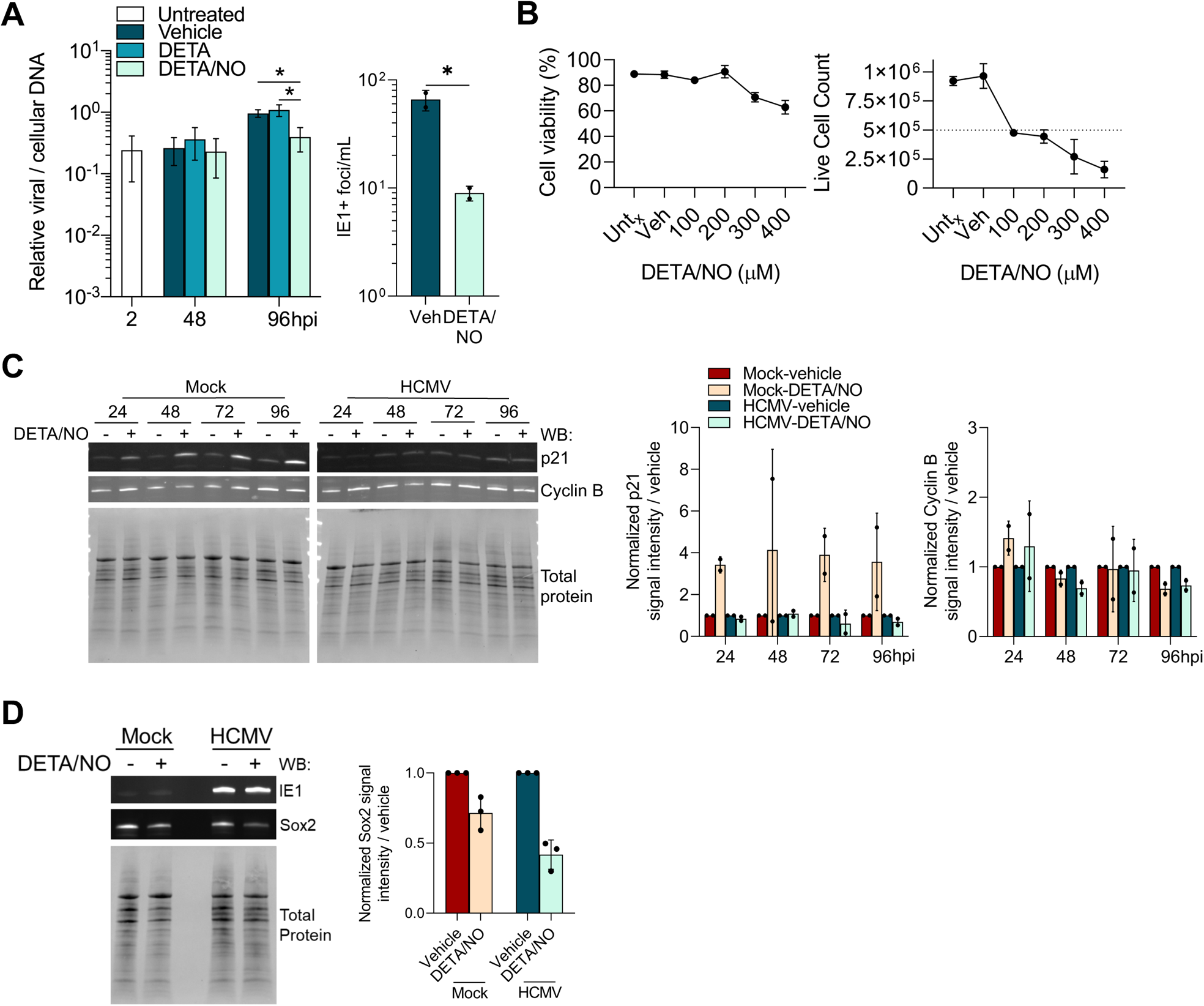
Nitric oxide attenuates HCMV replication while altering expression of proliferation and differentiation markers. **(A)** Neural progenitor cells (NPCs) were plated subconfluently and infected with HCMV strain TB40/E encoding free green fluorescent protein (TB40/E-GFP) propagated in ARPE-19 cells (TB40/EEpi-GFP) at an MOI of 3 infectious units/cell (IU/cell). Cells were treated at 2 hpi with 200 μM DETA/NO, DETA (spent donor), or vehicle control. Treatment was changed every 24 h. Relative viral to cellular DNA levels were determined at the indicated timepoints using primers to UL123 and TP53 genes. Viral titers determined by infectious units assay from culture supernatant collected at 96 hpi. Data are the result of two (viral titers) or three (DNA) biological replicates and two technical replicates. Errors bars represent SD. For viral DNA, statistics were performed on transformed data using a two-way ANOVA with multiple comparisons to determine significance. For viral titers, an unpaired, one-tailed t-test was used to determine significance. *P* < 0.05, *. **(B)** NPCs were plated subconfluently and treated every 24 h with the indicated concentrations of DETA/NO, vehicle control, or untreated (Untx). Cell viability and live cell number were quantified at 96 h post treatment using trypan blue exclusion. Data are the result of one-two biological replicates. Error bars represent SD. **(C and D)** NPCs were plated subconfluently and infected at an MOI of 3 IU/cell or mock-infected. Cells were treated at 2 hpi with 200 μM DETA/NO, or vehicle control and treatment was changed every 24 h. Whole cell lysates were collected, and protein levels were assessed by western blot using antibodies against the indicated proteins. Antibody signals were normalized to total protein before quantifying DETA/NO-treated relative to vehicle-treated for each timepoint using Image Lab software (Bio-Rad). Data represent two (D) or three (C) biological replicates.

Nitric oxide can have cytotoxic and cytostatic effects based on concentration. Therefore, we assessed these effects in uninfected NPCs. Cells were plated subconfluently at 5 x10^5^ cells/well and treated with a range of 100 to 400 μM DETA/NO, vehicle, or no treatment. Cell viability was quantified using trypan blue exclusion at 96 h post treatment. Cell viability at 100 and 200 μM concentrations were similar to vehicle and no treatment conditions (Fig. 4B). However, viability decreased at higher concentrations. We observed an initial increase in live cell number in untreated and vehicle-treated NPCs; however, this increase did not occur in 100 and 200 μM DETA/NO, suggesting nitric oxide induces cytostasis at specific concentrations. We observed a decrease in live cell count at 300 and 400 μM concentrations that was attributed to cell death (Fig. 4B). Overall, the results indicate that nitric oxide induces cytostasis and cell death in a concentration-dependent manner.

To investigate nitric oxide-induced cytostasis, we measured steady-state levels of cell cycle regulators cyclin-dependent kinase inhibitor p21^(CIP1/WAF1)^, which induces cell cycle arrest, and Cyclin B, which promotes cell cycle progression (Fig. 4C). Plated NPCs were infected as previously described at an MOI of 3 IU/cell or mock-infected and treated at 2 hpi with 200 μM DETA/NO or vehicle control. In uninfected NPCs, p21^(CIP1/WAF1)^ levels were increased during nitric oxide exposure by a respective 3.4-, 4.1-, 3.9-, and 3.6-fold at 24, 48, 72, and 96 hpi relative to vehicle. In contrast, levels remained unchanged in HCMV-infected cells. No changes were observed for Cyclin B levels. These data suggest that nitric oxide exposure induces p21^(CIP1/WAF1)^ expression, which likely contributes to cytostasis of uninfected NPCs.

### Nitric oxide decreases the transcription factor and multipotency regulator SOX2

Expression of p21^(CIP1/WAF1)^ and cell cycle exit are tightly linked with the onset of cellular differentiation (61, 62). We hypothesized that nitric oxide-induced cytostasis may interfere with NPC maintenance and differentiation in mock and HCMV-infected cultures. SOX2 is a master regulator required for maintaining multipotency and progenitor identity of NPCs (55–58, 63). Studies have demonstrated that SOX2 levels are reduced in an IE1-dependent manner during HCMV infection of embryonic-derived NPCs (13, 19). We initiated our studies by investigating SOX2 levels during nitric oxide exposure. At the same time, we investigated whether nitric oxide-mediated attenuation of HCMV replication could protect the cells from IE1-mediated downregulation of SOX2. Plated NPCs were mock or HCMV-infected as described above at an MOI of 3 IU/cell, treated at 2 hpi with 200 μM DETA/NO or vehicle control, and SOX2 levels determined at 96 hpi by western blot analysis (Fig. 4D). SOX2 levels were decreased by 0.3-fold relative to vehicle in uninfected NPCs. This reduction was more dramatic in HCMV-infected NPCs with a 0.6-fold decrease during nitric oxide exposure, suggesting the lack of a protective effect. Our data demonstrate that nitric oxide exposure reduces SOX2 levels and cellular proliferation and increases p21^(CIP1/WAF1)^ levels. Taken together, the results suggest that nitric oxide causes cell cycle arrest, suppresses progenitor cell maintenance, and influences early NPC differentiation.

### Mitochondrial function is compromised during nitric oxide exposure

Mitochondrial respiration is an important cellular function during differentiation when NPCs undergo a metabolic shift from mainly glycolytic to oxidative phosphorylation (64, 65). Nitric oxide can disrupt mitochondrial respiration by inhibiting complex I and IV of the electron transport chain (28–30). To determine if nitric oxide decreases mitochondrial respiration, we examined mitochondrial function using an extracellular flux assay (Fig. 5). NPCs were plated, incubated for 3 days, infected at an MOI of 3 IU/cell or mock-infected, and treated at 2 hpi with 200 μM DETA/NO, DETA (spent donor), or vehicle control. At 24 hpi, a mitochondrial stress assay was performed. Briefly, basal oxygen consumption rate (OCR) of cells is measured before sequential injections of oligomycin (ATP synthase inhibitor), carbonyl cyanide 4-(trifluoromethoxy)phenylhydrazone (FCCP, uncoupler of mitochondria), and rotenone/antimycin A (complex I and III inhibitors, respectively). This assay measures basal (green), ATP-linked (blue), and maximal (yellow) OCR (Fig. 5A). The difference of ATP-linked and maximal OCR is the spare capacity of the cells. A representative plot of the mitochondria stress assay is shown in Figure 5B with quantification of OCR shown in Figure 5C-F.

**Fig 5.**
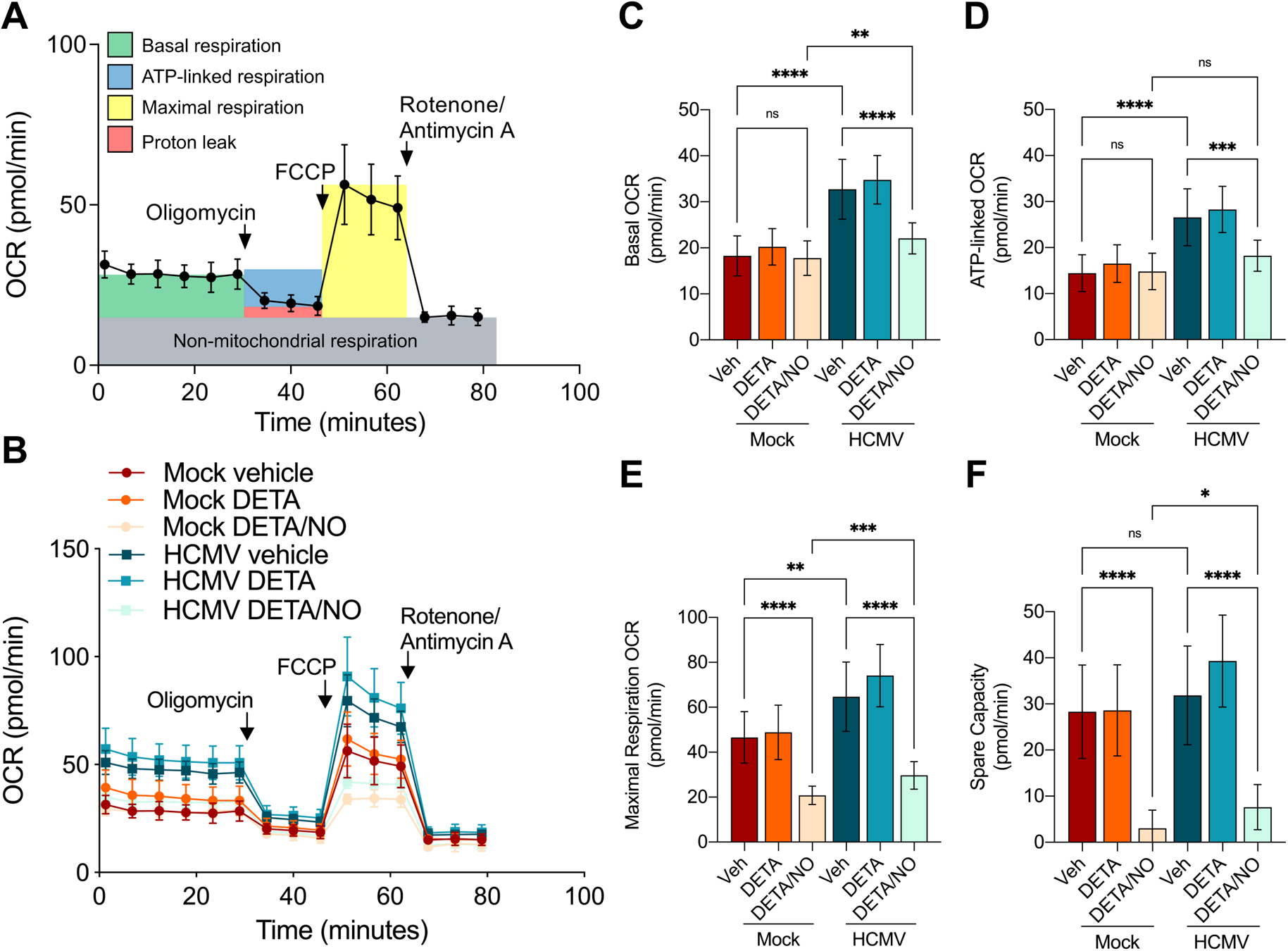
Mitochondrial function is compromised in mock and HCMV-infected NPCs during nitric oxide exposure. **(A)** Schematic of mitochondrial stress assay to assess mitochondrial function by measuring oxygen consumption rate (OCR). Sequential injections of oligomycin (1 uM), FCCP (1.5 μM), and rotenone/antimycin A (1 μM) allow the quantitation of basal (green), ATP-linked (blue), maximal (yellow), proton leak (red), and non-mitochondrial respiration (grey). **(B-F)** NPCs were plated at 50,000 cells/well in a Matrigel-coated 96-well Seahorse Bioscience dish and incubated for 3 d. Cells were infected with TB40/EEpi-GFP at an MOI of 3 IU/cell or mock-infected and treated at 2 hpi with 200 μM DETA/NO, DETA (spent donor), or vehicle control. Mitochondrial stress assay was performed at 1 dpi and media was replaced 1 h prior to assay with XF DMEM base medium without nitric oxide donor. Assay was run as described above, and basal **(C)**, ATP-linked **(D)**, maximal **(E)**, and spare capacity **(F)** OCRs were quantified. Data are the result of three biological replicates and five-six technical replicates. Error bars represent SD. Brown-Forsythe and Welch’s ANOVA tests with multiple comparisons were used to determine significance with DETA omitted from analysis. *P* = 0.03, *; *P* < 0.007, **; *P* < 0.0002, ***; *P* < 0.0001, ****; ns = not significant.

Basal OCR of HCMV-infected cells was significantly increased by 79% compared to uninfected NPCs (Fig. 5C), indicating infection increases mitochondrial respiration by 24 hpi in NPCs. However, nitric oxide exposure reduced basal OCR by 33% in infected NPCs but not in uninfected cells. ATP-linked OCR also increased by 84% in vehicle-treated, HCMV-infected cells compared to vehicle-treated, uninfected NPCs (Fig. 5D). This rate was reduced by 31% upon nitric oxide exposure of HCMV-infected cells. No differences were observed in mock-infected conditions. Maximal respiration of NPCs was increased by 39% during infection (Fig. 5E), indicating an increase in the maximum range of the cells’ response to energy demands. However, in contrast to basal and ATP-linked OCR, maximal respiration was reduced in both uninfected and HCMV-infected NPCs by a respective 55% and 54% during exposure to nitric oxide. Likewise, the spare respiratory capacity of the cells was reduced by a respective 89% and 76% in mock and HCMV-infected conditions during nitric oxide exposure (Fig. 5F). OCRs of the spent donor control (DETA) did not deviate from those of vehicle for the uninfected or HCMV-infected groups and were not included in the statistical analysis (Fig. 5C-F). These data demonstrate that nitric oxide significantly suppresses HCMV-mediated increase of basal and ATP-linked respiration. Further, nitric oxide reduces the capacity of NPCs to respond to energy demands through mitochondrial respiration regardless of infection.

### Nitric oxide limits neuron and glial differentiation

Reduced mitochondrial respiration, decreased SOX2 expression, and cytostatis led us to hypothesize that nitric oxide restricts neuron and glial cell differentiation from NPCs. To test this hypothesis, we investigated the effect of nitric oxide and HCMV on neural differentiation at 14 days post differentiation. Cells were cultured in the absence of growth factors to allow spontaneous differentiation (16, 59). We treated with 200 μM DETA/NO every 48 h to reduce the chance of cytotoxicity due to prolonged exposure and reduced the MOI to avoid loss of cultures due to lytic replication. We modeled steady-state nitric oxide levels released from 200 μM DETA/NO and estimated levels decrease from approximately 1.25 to 0.5 μM over 48 h (Fig. 6A). These levels are in contrast to treatment every 24 h that range from 1.25 to 1 μM (Fig. 6A). Following the experimental timeline in Figure 6B, NPCs were plated for 3 days and infected at an MOI of 0.05 IU/cell or mock-infected. Subsequently, cells were treated at 2 hpi and every 48 h with DETA/NO, DETA (spent donor), or vehicle control out to 11 dpi. We assessed markers of neural populations using immunofluorescence. We used the marker Nestin to identify the progenitor population as SOX2 is reduced during HCMV infection of embryonic stem cell-derived NPCs (13). Further, we identified neuron populations by labeling for Tuj1, a class III beta-tubulin specific to neurons, and glial populations by labeling for glial fibrillary acidic protein (GFAP), a type III intermediate filament protein expressed by glial cells (**Fig. S1A**). In mock, vehicle-treated cultures, we observed expression of each developmental marker, indicating a mix of neural populations. We observed Nestin-positive cells contained small processes with large cell bodies (**Fig. S1A**), while Tuj1-positive cells had long, netted neurites with small cell bodies (**Fig. S1A**). GFAP-positive cells had long projections with a large cell body (**Fig. S1A**). HCMV-infected cultures contained single nuclei and multi-nucleated GFP-positive (GFP+) cell bodies (**Fig. S1B**), indicating syncytia formation regardless of nitric oxide exposure. We attempted to quantify progenitor, neuron, and glial populations using immunofluorescence since our goal was to evaluate changes in marker expression. However, we were unable to use this approach due to culture density and overlapping projections (**Fig. S1A**).

**Fig 6.**
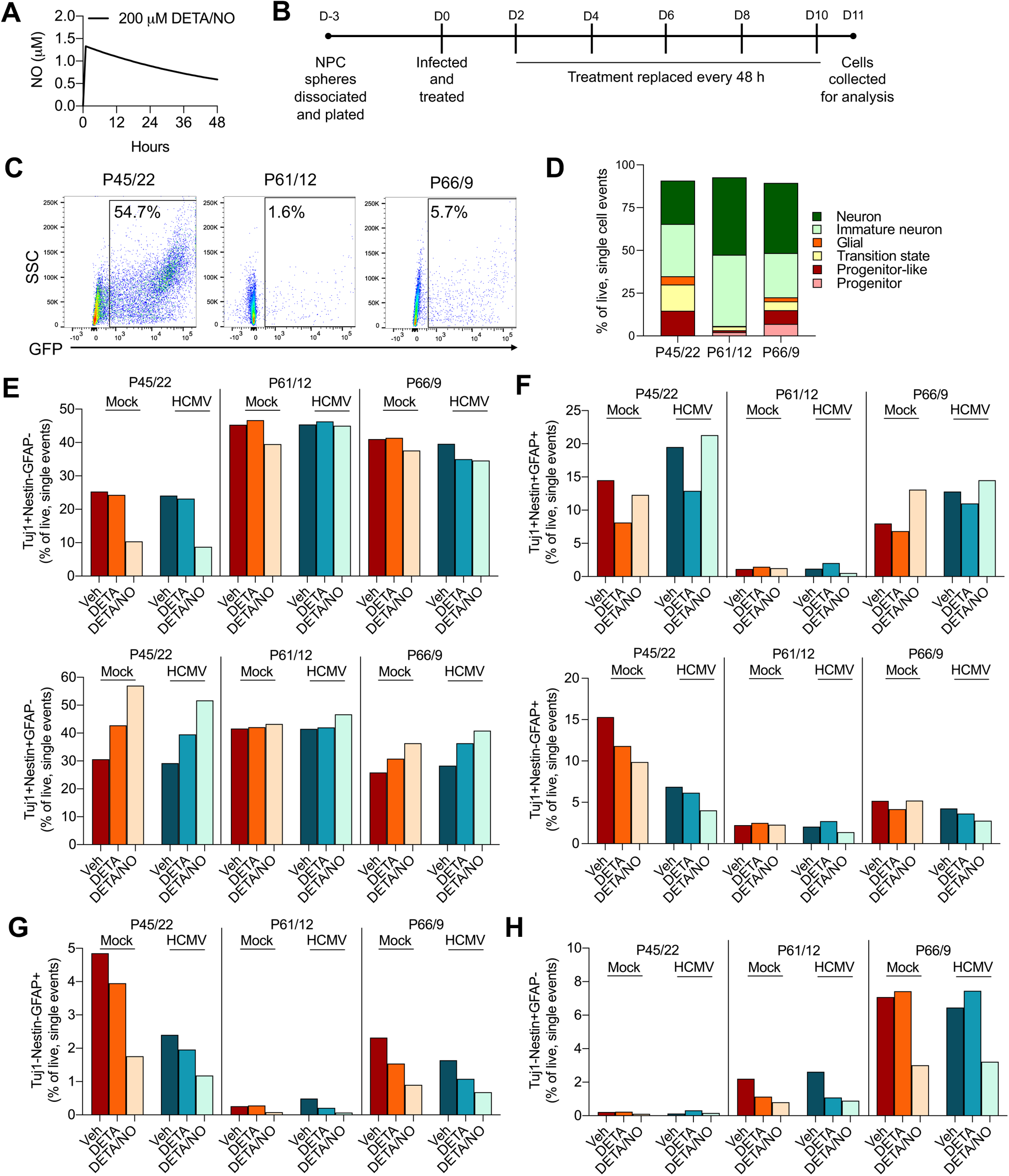
Nitric oxide alters NPC differentiation but is impacted by cell passage. **(A)** Steady state nitric oxide levels for 200 μM DETA/NO were estimated using COPASI software (96). The rate of decay for DETA/NO was calculated from the *t1/2* 20 h at 37°C. The estimated oxygen concentration was set to 220 μM. The reaction rate of nitric oxide with oxygen to form nitrogen dioxide (NO2) was entered as 2.4 x 10^6^ M^-2^ S^-1^ (97), and the reaction rate of nitric oxide with NO2 to form nitrite was set to 1 x 10^9^ M^-2^ S^-1^. **(B)** Experimental timeline for NPC differentiation. NPCs were plated subconfluently and infected with TB40/EEpi-GFP at an MOI of 0.05 IU/cell or mock-infected. Cells were treated at 2 hpi with 200 μM DETA/NO, DETA (spent donor) or vehicle control, and treatment was changed every 48 h. At 11 dpi, NPC differentiation was assessed using various methods. **(C-H)** Following the experimental timeline defined in A, progenitor, glial, and neuron populations were identified by labeling for Nestin, GFAP, and Tuj1, respectively, and quantified using flow cytometry at 11 dpi. Data are from three different cell passages with passage number indicating iPSC to NPC neurosphere passage. (C) HCMV-infected cell populations were identified using GFP expression as an indicator of infected cells. Live, single cell events were gated and GFP-negative (GFP-) and GFP-positive (GFP+) gates were applied to the unstained, uninfected cells. These gates were then applied to the experimental samples and percent GFP+ relative to live, single cell events is displayed. **(D-H)** Neural markers were quantified by gating on live, single cell events and using fluorescent minus one controls to gate Tuj1/GFAP, Tuj1/Nestin, and GFAP/Nestin populations. **(D)** Frequency variability between cell passage for populations of Tuj1+Nestin-GFAP-(neuron), Tuj1+Nestin+GFAP-(immature neuron), Tuj1-Nestin-GFAP+ (glial), Tuj1+Nestin-GFAP+ (transition state), Tuj1+Nestin+GFAP+ (progenitor-like), and Tuj1-Nestin+GFAP-(progenitor) populations. **(E)** Tuj1+Nestin-GFAP- and Tuj1+Nestin+GFAP-populations were identified by gating live, single cell events and gating for Tuj1/GFAP populations. Tuj1+GFAP-events were then gated for Tuj1/Nestin populations. **(F)** Tuj1+Nestin+GFAP+ and Tuj+Nestin-GFAP+ were identified following a similar workflow but Tuj1+GFAP+ events were gated for Tuj1/Nestin. **(G)** Tuj1-Nestin-GFAP+ populations were identified by gating live, single cell events and gating for Tuj1/GFAP populations. Tuj1-GFAP+ events were then gated for GFAP/Nestin populations. **(H)** Tuj1-Nestin+GFAP-were identified by gating for live, single cells and Tuj/GFAP population and gating for Nestin on the Tuj1-GFAP-population. All populations were quantified as percent of live, single cell events.

To overcome this limitation, we switched to flow cytometry to quantify differentiated neural populations, which also allowed us to identify double- and triple-marker positive cells (66). As previously described, NPCs were plated from dissociated iPSC-derived NPC neurospheres and completed using different NPC passages. The passage numbers are recorded as iPSC to NPC neurosphere passage number (iPSC/Neurosphere). Cells from passage 45/22, 61/12, and 66/9 were used for these studies. NPCs were mock-infected or infected as outlined in Fig. 6B, and flow cytometry performed at 11 dpi. Cells were labeled with Nestin, Tuj1, and GFAP-conjugated fluorescent antibodies. Gates were placed to isolate live, single cell events, and GFP+ events were identified by first gating on the unstained, uninfected population (**Fig. S2A**). Forward and side scatter gates were extended to include the larger GFP+ cells. It is possible that syncytia were lost despite the increased gate size; however, we reasoned that these cell bodies would be equally lost between conditions. Uninfected cells had no GFP+ events. GFP+ events varied in infected cultures depending on the passage of NPCs, despite infecting at the same MOI (Fig. 6C). For cells from passage 45/22 (P45/22), 54.7% of the live, single cell events were GFP+. In P61/12 and P66/9 cells, GFP+ events were 1.6 and 5.7%, respectively. As GFP levels varied between experiments, we grouped GFP+ and GFP-negative (GFP-) events to analyze the whole culture population for each experiment. We used fluorescence minus one (FMO) controls (**Fig. S2B)** to define Tuj1/GFAP, Tuj1/Nestin, and GFAP/Nestin populations, and our gating strategy is shown in **Figure S2C.**

We first determined frequencies of Tuj1+Nestin-GFAP-events, which represent neuron populations, and Tuj1+Nestin+GFAP-events, which likely represents an immature neuron population as they express both Tuj1 and Nestin. We quantified these populations by gating on live, single cell, Tuj1+ events from Tuj1/GFAP populations (**Fig. S2C,** Q1). Tuj1+Nestin-GFAP-(neurons) frequencies in the mock, vehicle-treated control groups varied between passage number. For P45/22 cultures, the frequency of Tuj1+Nestin-GFAP-(neurons) events was 25% compared with the respective 45% and 41% quantified in P61/12 and P66/9 cultures (Fig. 6D & E). This variation indicates intrinsic differences in the spontaneous differentiation capacity of the NPCs. We suspect these differences also contribute to the variability in susceptibility to infection (Fig. 6C). Upon exposure to nitric oxide, we observed a substantial decrease in the percent of Tuj1+Nestin-GFAP-(neurons) events in P45/22 cultures in mock and HCMV-infected populations (Fig. 6E). Less dramatic reductions were observed in P61/12 and P66/9 (Fig. 6E), suggesting that sensitivity to nitric oxide is influenced by intrinsic differences between passages and developmental states. The percent of Tuj1+Nestin+GFAP-(immature neurons) events for mock, vehicle-treated cells were more similar between P45/22 and P66/9 at a respective 31% and 26% (Fig. 6D & E). This population was increased in P45/22 and P66/9 cultures during nitric oxide exposure regardless of infection (Fig. 6E). For P61/12, the percent of Tuj1+Nestin+GFAP-(immature neurons) events for mock, vehicle-treated cultures was 42% and no differences were observed during nitric oxide exposure (Fig. 6E). Again, these data suggest that differentiation capacity between cell passages influences their susceptibility to nitric oxide. Cells exposed to the spent donor (DETA) had similar frequencies of Tuj1+Nestin-GFAP-(neurons). For P45/22 and P66/9, there was an increased percent of Tuj1+Nestin+GFAP-(immature neurons) compared to vehicle though not to the level of nitric oxide exposed cells (Fig. 6E). This result suggests a limited impact of nitric oxide oxidation products on neuron differentiation. Overall, the results suggest that nitric oxide reduces neuron populations.

We next examined percent of Tuj1+Nestin+GFAP+ and Tuj1+Nestin-GFAP+ (Fig. 6F) relative to live, single cells by gating on Tuj1+GFAP+ events (**Fig. S2C**, Q2). Tuj1+Nestin+GFAP+ populations likely represent progenitor-like cells (67, 68). The identity of Tuj1+Nestin-GFAP+ populations is unknown but may represent cells in transition to a neuron or glial state after downregulation of Nestin (67). Frequencies of Tuj1+Nestin+GFAP+ (progenitor-like) events varied between passages for mock, vehicle-treated cultures with levels most similar between P45/22 and P66/9 (Fig. 6D & F). We observed an increase in the P66/9 progenitor-like population in uninfected nitric oxide exposed cultures that was not observed in other passages (Fig. 6D & F). P61/12 had substantially less Tuj1+Nestin+GFAP+ (progenitor-like) events in mock, vehicle-treated cultures (Fig. 6D & F). DETA treatment decreased frequencies of Tuj1+Nestin+GFAP+ events in P45/22 and P66/9, suggesting some impact of nitric oxide oxidation products on this progenitor-like population (Fig. 6F). Tuj1+Nestin-GFAP+ (transition state) populations were also variable between passages in mock, vehicle-treated cultures at 15.3%, 2.2%, and 5.2% for P45/22, P61/12, and P66/9, respectively (Fig. 6D & F). This population was reduced during infection of P45/22, and nitric oxide exposure also reduced these frequencies regardless of infection (Fig. 6F). Overall, our data indicate a limited impact of nitric oxide on Tuj1+Nestin+GFAP+ (progenitor-like) and Tuj1+Nestin-GFAP+ (transition state) populations.

Percent of Tuj1-Nestin-GFAP+ cells, representing glial populations, were obtained by gating on Tuj1-GFAP+ (**Fig. S2C**, Q3) and then gating for Tuj1-Nestin-GFAP+ (Fig. 6G). We again observed differences in the frequency of glial populations between the passages of mock, vehicle-treated groups with a respective 4.8%, 0.2%, and 2.3% for P45/22, P61/12, and P66/9 (Fig. 6D & G). For P45/22 and P66/9, Tuj1-Nestin-GFAP+ (glial) populations for infected, vehicle-treated cultures were modestly decreased compared with mock, vehicle-treated. We observed a limited impact from the spent donor on this population (Fig. 6G), suggesting oxidation products do not alter glial populations. Nitric oxide decreased Tuj1-Nestin-GFAP+ (glial) frequencies regardless of infection (Fig. 6G), suggesting nitric oxide limits glial differentiation. Frequencies of Tuj1-Nestin+GFAP+, which may represent immature glial cells, were below 1% and not included in further analysis (data not shown).

Finally, levels of Tuj1-Nestin+GFAP-, representing progenitor populations were obtained by gating on Tuj1-GFAP-events and Nestin+ events (**Fig. S2C**, Q4). Levels of Tuj1-Nestin+GFAP-(progenitor) were notably higher in P66/9 compared to P45/22 and P61/12 (Fig. 6D & H). Nitric oxide decreased progenitor populations in P66/9 mock, vehicle-treated cultures from 7.1% to 3% with a similar reduction observed in infected cultures (Fig. 6H). The frequencies of progenitor populations for P45/22 and P61/12 were too low to observe differences (Fig. 6H). Taken together, our data suggest nitric oxide, and to some degree oxidation products, disrupts terminal differentiation of both neuron (Tuj1+Nestin-GFAP-) and glial (Tuj1-Nestin-GFAP+) populations with a trend toward increased immature neurons (Tuj1+Nestin+GFAP-) regardless of infection.

### Despite HCMV inhibition, nitric oxide fails to rescue HCMV-mediated defects in purinergic receptor and voltage gated calcium channel activity

Calcium signaling through purinergic receptors in glial cells and voltage gated calcium channels (VGCC) in neurons is a critical process in CNS development (69–71). We have previously demonstrated that HCMV-infected differentiated neural cultures have a loss of purinergic and VGCC Ca^2+^ signaling (20). Because we observed a decrease in both neuron and glial populations, we hypothesized that nitric oxide exposure may result in a loss of calcium response. Replicates were completed using NPCs from P61/9, 61/10, 61/12, and 61/14. Cells were plated, incubated for 3 days, and infected at an MOI of 0.05 IU/ cell or mock-infected. At 2 hpi, cells were treated with DETA/NO, DETA, or vehicle control and treatment changed every 48 h (Fig. 6B). At 11 dpi, cells were subjected to live cell Ca^2+^ imaging (Fig. 7). Briefly, cultures were incubated in Fura-2-acetoxymethyl ester (Fura-2AM), a ratiometric Ca^2+^ sensitive dye. This dye excites at 340 nm and 380 nm in the Ca^2+^-bound and unbound state, respectively, and is measured as the 340/380 nm ratio. Cells were stimulated with either 10 nM ATP for purinergic receptors or 50 nM KCl for VGCC and washed between conditions. Baseline levels were measured 30 s before stimulation, and cells were considered responding if the 340/380 nm ratio increased 5% over baseline. We quantified baselines before stimulation, percent of cells responding to stimulation, and maximum amplitude of those cells for mock and infected cultures. HCMV-infected GFP+ and GFP-cells from the same culture were analyzed separately (Fig. 7). Consistent with our previous studies (20), GFP+ cells had decreased baseline, percent of cells responding, and maximum amplitude compared to mock and GFP-groups. However, these comparisons were not included in the statistical analysis. For the different conditions, we observed no significant differences in baseline prior to stimulation with ATP (Fig. 7A) or KCl (Fig. 7B) for any of the groups, indicating neither nitric oxide nor spent donor impact baseline Ca^2+^ levels. The percent of cells responding to ATP stimulation in the presence of DETA and DETA/NO were increased by 3.6- and 3.0-fold for mock and GFP-groups compared to vehicle; however, only mock, DETA-treated groups were significantly different (Fig. 7A). Similarly, the percent of cells responding to KCl stimulation in the presence of DETA and nitric oxide were increased by a respective 2.0- and 1.4-fold compared to vehicle control but only GFP-, DETA-treated groups were significantly different (Fig. 7A). Together, these results indicate that nitric oxide oxidation products contribute to the Ca^2+^ response to ATP and KCl stimulation. Finally, the maximum amplitude of the response to ATP, but not KCl, was significantly increased in mock-infected cells upon exposure to DETA and DETA/NO by 2.3- and 3.3-fold, respectively (Fig. 7A). This response was also elevated in GFP-DETA and nitric oxide exposed groups by a respective 5.2- and 4.3-fold (Fig. 7A). Together, our data indicate that long-term nitric oxide exposure, including oxidation products, increases the number of cells responding to ATP and KCl stimulation with increased amplitude of response to ATP. Further, despite this response and inhibition of HCMV, nitric oxide does not rescue HCMV-mediated disruption of Ca^2+^ responses.

**Fig 7.**
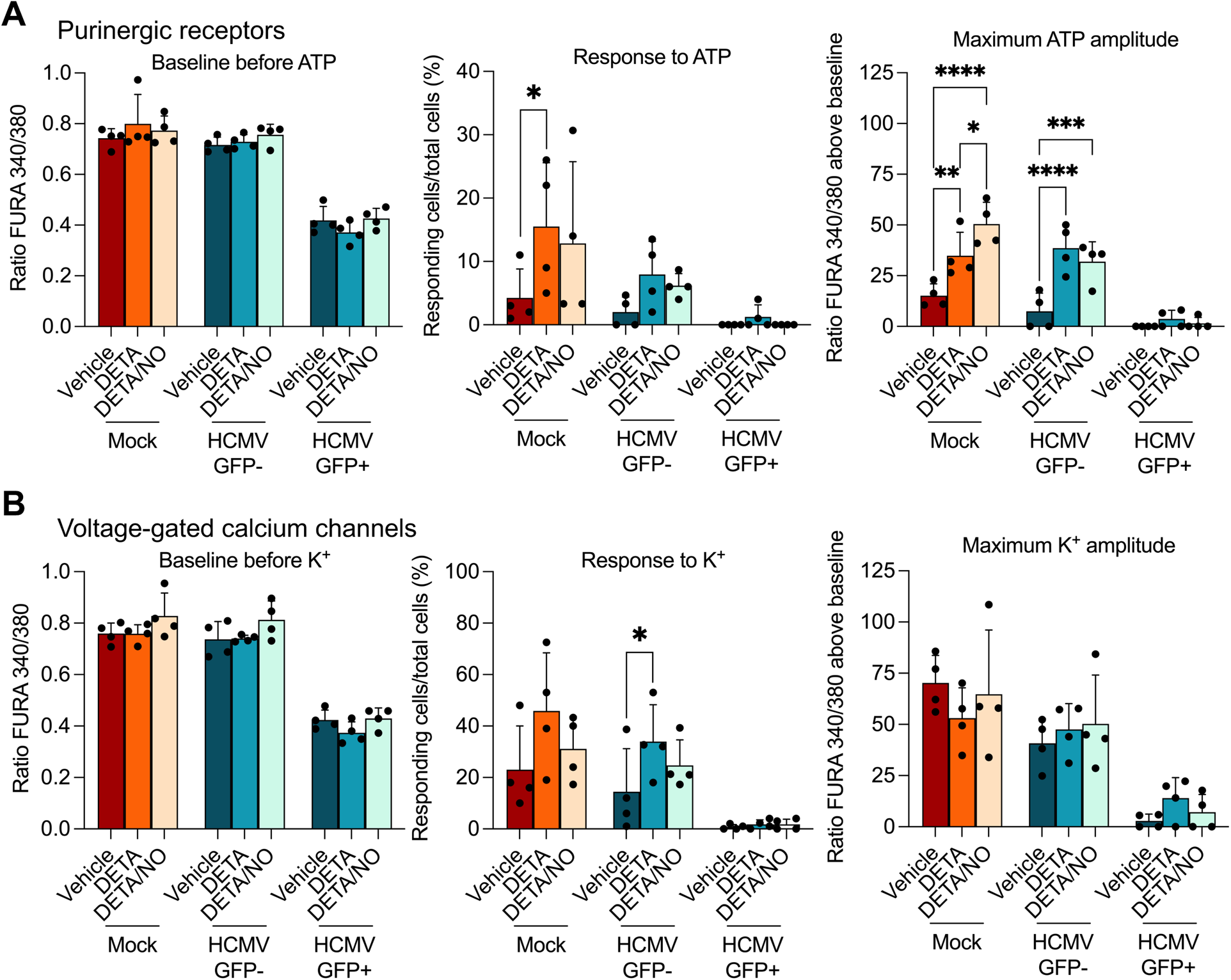
Nitric oxide increases amplitude of cellular calcium response to ATP but not K+ and does not rescue HCMV-mediated defects in purinergic and VGCC receptor activity. Following the experimental timeline defined in Fig 6B, cells were analyzed using live-cell Ca^2+^ imaging with FURA 2AM. **(A)** Intracellular Ca^2+^ levels (baseline) for mock, HCMV-infected GFP-negative (GFP-) and GFP-positive (GFP+) were measure prior to stimulation of purinergic receptors with ATP (left panel) after removed of nitric oxide donor. Cells responding to 10 μM ATP were counted and quantified as percent of total cells (middle panel). Maximum amplitude of ATP response was quantified from the cells that displayed a FURA 340/380 ratio increase of 5% or more above baseline to 10 μM ATP (right panel). **(B)** Intracellular Ca^2+^ levels (baseline) for mock, HCMV-infected GFP- and GFP+ were measure prior to stimulation of VGCC with KCl (left panel) and removal of nitric oxide donor. Cells responding to 50 μM KCl were counted and quantified as percent of total cells (middle panel). Maximum amplitude of KCl response was quantified from the cells that displayed a FURA 340/380 ratio increase of 5% or more above baseline to 50 μM KCl. Data are the result of four biological replicates. Statistics were performed on transformed data for response to ATP and K+. A two-way ANOVA with multiple comparisons was used to determine significance. *P* ≤ 0.04, *; *P* = 0.007, **; *P* < 0.0008, ***; *P* < 0.0001, ****.

## Discussion

Nitric oxide is a selectively reactive free radical produced as a component of the innate immune response and has antiviral activity against HCMV replication. Prior to our studies, the impact of nitric oxide on the developing brain during HCMV infection was unknown. We modeled this complex relationship using 3-dimensional cortical organoids that recapitulate many aspects of fetal brain development. We observed that nitric oxide disrupts HCMV replication. However, nitric oxide substantially disrupts tissue organization and neural rosette structure in both mock and HCMV-infected organoids (Fig. 2 and **3**). Gabrielli et al. (7) observed that brains from congenitally-infected fetuses have numerous necrotic brain regions with infiltrating immune cells that include macrophages and microglia that are known to produce nitric oxide (24, 25). These observations suggest that nitric oxide production during congenital infection is likely highest at the sites of infection with surrounding uninfected cells exposed to a gradient of nitric oxide. It is likely that as the nitric oxide gradient decreases there are less pathogenic consequences on uninfected cells. In our studies, the outer layer of the organoid is exposed to the highest concentration of nitric oxide, which does not fully mimic conditions during infection. To better recapitulate physiological conditions, cortical organoids can be differentiated to include microglia, which produce nitric oxide via NOS2 during infection (25, 72). Studies by Ormel et al (72) demonstrated that organoids are capable of producing mesodermal progenitors that differentiate to cells expressing classical microglia markers. The addition of microglia to the cortical organoid model of HCMV infection will allow investigation into the role of other immune molecules such as cytokines on developing cortical tissue.

Microcephaly, or smaller head and/or brain size, is associated with disruption of progenitor populations (54). HCMV infection can result in microcephaly, which contributes to cognitive defects in congenitally-infected infants (2). Dysregulation of progenitor populations during HCMV infection is thought to contribute to this birth defect (20, 22, 23). Our studies suggest that nitric oxide may also contribute to microcephaly as exposure decreased SOX2, the regulator of progenitor maintenance, and altered neural populations, including decreased neurons (Tuj1+Nestin-GFAP-) and glial (Tuj1-Nestin-GFAP+) cells (Fig. 5E-H). Further, cortical organoids exposed to nitric oxide had significant disruption of tissue organization and rosette structures, suggesting dysregulated progenitor populations (Fig. 2 and **3**).

Metabolic reprogramming is essential for neuronal differentiation (65, 73–75). Agostini et al. (73) and Zheng et al. (65) demonstrated that basal and maximal OCR is increased in differentiated neurons compared to progenitor cells. Zheng et al. (65) also showed that a reduction in glycolytic enzymes is essential for neuron differentiation. Our data demonstrate that nitric oxide significantly decreases maximal respiration of NPCs and reduces neuron (Tuj1+Nestin-GFAP-) populations during NPC differentiation in both uninfected and HCMV-infected cultures (**Fig. 6E, 6F,** and **5E**). It is possible that nitric oxide limits the potential of NPCs to differentiate by limiting the capacity of the cell to respond to increased energy demands and forcing glycolysis to compensate for limited energy. Further, dysregulation of metabolism by HCMV infection is a plausible mechanism for the reduction in neuron populations observed during HCMV infection (13, 16). It is also possible that nitric oxide-mediated cell death is resulting in decreased mature neuron populations. Nitric oxide can induce neuron death through several mechanisms, including inhibition of mitochondrial respiration, altered metabolism, poly-ADP-ribose polymerase (PARP) activation, and/or formation of reactive nitrogen species (76). Future studies will elucidate the mechanisms of limited neural differentiation versus cell death during HCMV infection and nitric oxide exposure.

Ca^2+^ signaling is crucial for various stages of neural development (77, 78). Our previous work demonstrated that HCMV infection reduced calcium responses of dissociated organoids and NPCs (20), and Sun et al (23) demonstrated disrupted calcium response in whole cortical organoids. Our data indicate that long-term nitric oxide exposure does not rescue HCMV-mediated defects in Ca^2+^signaling despite evidence of attenuated viral replication (Fig. 7 and **4A**). Our data suggest that loss of Ca^2+^ response during HCMV infection does not require active replication and/or that the level of HCMV attenuation by nitric oxide is not sufficient to rescue. These results are consistent with previous studies in fibroblasts that indicate Ca^2+^ dysregulation can occur at various stages of infection, including entry and early replication events (79–82).

Mitochondrial respiration generates ATP through the proton gradient established by the electron transport chain (ETC). The electron donors NADH and FADH2 are critical contributors to mitochondrial respiration as they donate electrons to complex I and II, respectively, of the ETC. HCMV infection increases mitochondrial biosynthesis, and functional mitochondria are required for efficient viral replication (83, 84). Our previous study determined that nitric oxide inhibits HCMV replication in fibroblasts and epithelial cells by a multi-factorial mechanism that includes altered metabolism and decreased mitochondrial respiration during infection (48). Here, we demonstrate that nitric oxide also inhibits HCMV spread in cortical organoids and HCMV replication in NPCs (Fig. 1 and **4**). Nitric oxide may have a similar mechanism of inhibition in NPCs as other cell types. In HCMV-infected fibroblasts, basal OCR is increased at 48 hpi but not 24 hpi (48, 84). In contrast, we observed an increase in basal OCR at 24 hpi in NPCs (Fig. 6C), suggesting differences in mitochondrial and metabolic manipulation in NPCs during infection. Nitric oxide reduced basal respiration in HCMV-infected NPCs to the level of uninfected cells. Direct inhibition of ETC enzymes could decrease basal respiration (28, 85); however, this mechanism is unlikely as there is no effect on basal levels by nitric oxide in uninfected NPCs (Fig. 6C). More likely, nitric oxide alters metabolic intermediates that are increased during infection and required for the ETC such as NADH and FADH2 from the TCA cycle. Chambers et al. (86) and others demonstrated that glutamine uptake and glutaminolysis are essential for infection by providing intermediates to the TCA cycle (87–89), and our previous studies suggest that glutamine diversion has a role in nitric oxide-mediated inhibition of HCMV in fibroblasts (48). If nitric oxide is diverting glutamine from its primary role in NPCs during HCMV infection, it could account for decreased basal respiration, and ultimately, viral replication. Future studies will elucidate the metabolic impact on HCMV-infected NPCs during nitric oxide exposure.

In summary, congenital HCMV infection and its impact on the developing fetal brain remain a serious health concern. Using a cortical organoid model to recapitulate infection of the fetal brain, we have demonstrated that the free radical nitric oxide contributes to developmental defects during HCMV infection despite its antiviral activity. Further, this study provides additional evidence that indirect effects of HCMV infection such as nitric oxide production are potent contributors to disease.

## Materials and Methods

### Cell culture and virus

Undifferentiated induced pluripotent stem cell (iPSC) colonies, initially derived from fibroblasts (Coriell fibroblast line GM003814) (52, 53), from a healthy individual were maintained in Essential 8 medium (ThermoFisher Scientific) and grown under feeder-free conditions on Matrigel (Corning). Neural progenitor cells (NPCs) were differentiated and maintained as neurospheres (59) in Stemline (Millipore Sigma) supplemented with 0.5% N-2 supplement (ThermoFisher Scientific), 100 ng/ml EGF (Miltenyi Biotech), 100 ng/ml fibroblast growth factor (FGF; Stem Cell Technologies), and 5 μg/ml heparin (Millipore Sigma). Neurospheres were dissociated using TrypLE and plated at 500,000 to 600,000 cells per well on Matrigel-coated dishes. Plated NPCs were differentiated in Neurobasal medium (ThermoFisher Scientific) supplemented with 2% B-27 (ThermoFisher Scientific).

TB40/EFb-eGFP stocks were obtained by transfecting a bacterial artificial chromosome (BAC) encoding HCMV strain TB40/E-eGFP and a plasmid encoding UL82 into MRC-5 fibroblast using electroporation at 260 mV for 30 ms with a 4-mm-gap cuvette and a Gene Pulser Xcell electroporation system (90–93). TB40/EEpi-eGFP stocks were obtain by infecting ARPE-19 cells with TB40/EFb-eGFP. Viral media was collected and pelleted through a sorbitol (20% sorbitol, 50 mM Tris-HCl, pH 7.2, 1 mM MgCl2) cushion at 20,000 x g for 1 h in a Sorvall WX-90 ultracentrifuge and SureSpin 630 rotor (Thermo Fisher Scientific). Viral stock titers were titered on MRC-5 (TB40/EFb-eGFP) or ARPE-19 (TB40/EEpi-eGFP) cells in 96-well dishes using a limiting dilution assay (TCID50). GFP-positive wells were determined at 2 weeks post infection and resulting titers were referred to as infectious units per milliliter (IU/mL). MOI of 0.05 or 3 infectious units per cell (IU/cell) were used for infection of NPCs using TB40/EEpi-eGFP. Day 35 cortical organoids were infected using an MOI of 500 infectious units per microgram using TB40/EFb-eGFP.

### Cortical organoids

Cortical organoid cultures were differentiated from the healthy iPSC cell line according to the specification of the cerebral organoid kit from StemCell Technologies (#08570) that relies on an established protocol (54). Briefly, iPSCs were plated at 9,000 cells per well onto 96-well ultralow attachment plates for embryoid body (EB) formation and grown in EB formation media (StemCell Technologies). At day 5, the induction of neural epithelium was initiated by moving the EBs into an ultralow attachment 24-well plate and feeding with induction media (StemCell Technologies). On day 7, neural tissues were embedded in Matrigel droplets and moved to ultralow attachment 6-well plates and fed expansion media (Stemcell Technologies). Day 10 organoids were transferred to a rocker to elicit the circulation of nutrients and prevent organoids from sticking to the dish. The organoids were expanded and fed with fresh maturation media every 3-4 days. This method is also described in (20).

### Chemical reagents

Stocks of 50 mM diethylenetriamine NONOate (DETA/NO) (Cayman Chemicals) in 0.01 M NaOH were maintained at −20°C. Spent donor (DETA) was prepared by diluting DETA/NO to the appropriate concentration in medium and incubating at 37°C for approximately 72 h and then maintained at 4°C. DETA/NO cytotoxicity was assessed by plating 500,000 NPCs/well on Matrigel (Corning) coated 6-well dishes and treated with a range of concentrations or vehicle control (NaOH). Treatment was replaced every 24 h and cell viability and number were quantified at 96 h post treatment using a hemocytometer and trypan blue exclusion. For the remaining experiments, 200 μM and 400 μM DETA/NO were used to treat NPCs and cortical organoids, respectively.

### Analysis of nucleic acid and protein

Quantitative PCR (qPCR) was used to quantify viral DNA levels. Cells were collected by trypsinization and isolated using the DNeasy blood and tissue kit (Qiagen). Primers to HCMV UL123 (5′-GCCTTCCCTAAGACCACCAAT-3′ and 5′-ATTTTCTGGGCATAAGCCATAATC-3′) and cellular TP53 (5′-TGTTCAAGACAGAAGGGCCTGACT-3′ and 5′-AAAGCAAATGGAAGTCCTGGGTGC-3′) (IDT) were used for quantitation. qPCR was completed using SYBR green PCR mix (ThermoFisher) and QuantStudio 6 Flex real-time PCR. Relative HCMV UL123 was normalized to cellular TP53.

Protein levels were determined by western blot analysis. Cells were collected by trypsinization, resuspended in lysis buffer (50 mM Tris-HCl, pH 8.0, 150 mM NaCl, 1% SDS) with protease and phosphatase inhibitor, and lysed by sonication. Protein concentrations were determined by Pierce Bicinchoninic Acid (BCA) assay kit (ThermoFisher) and 20 ug of protein were resolved by SDS-PAGE using 4-15% gradient gels. Proteins were transferred to nitrocellulose membrane (GE Healthcare Life Sciences) using a semidry Trans-Blot transfer system (Bio-Rad), and membranes blocked for 1 h in 5% milk in PBS-T (phosphate-buffered saline, 0.1% Tween 20) or TBS-T (tris-buffered saline, 0.1% Tween 20). After blocking, membranes were incubated with primary antibody diluted in 5% milk or bovine serum albumin (BSA) in PBS-T or TBS-T overnight at 4°C. Membranes were washed in PBS-T or TBS-T and incubated with StarBright Blue secondary antibody (Bio-Rad) diluted in 5% milk in PBS-T for 30 min at room temperature in the dark and imaged using a Bio-Rad ChemiDoc imager.

The following antibodies and dilutions were used for western blot analysis: rabbit anti-cyclin B1 (clone D5C10, 1:1,000; CST), mouse anti-p21 (clones CP36 and CP74, 1:1,000; Millipore Sigma), and rabbit anti-SOX2 (1:1000; Millipore Sigma). The HCMV IE1 (clone 1B12, 1:1000) antibody was generously provided by Tom Shenk (Princeton University). Secondary antibody of StarBright Blue goat anti-mouse or anti-rabbit (Bio-Rad) were used at 1:10,000 dilution.

### Mitochondrial Stress Assay

A Seahorse XFe96 Analyzer was used to determine oxygen consumption rates (OCR). For NPCs, 50,000 cells/well were plated on Matrigel (Corning) coated Agilent Seahorse 96-well dishes and incubated for 3 d. Cells were infected with HCMV and treated as described in the figure legend. The mitochondrial stress analysis was performed at 1 dpi with media replaced 1 h prior with XF DMEM base medium. The assay was performed by sequential injections of oligomycin (1 μM), carbonyl cyanide 4-(trifluoromethoxy)phenylhydrazone (FCCP) (1.5 μM), and rotenone/antimycin A (1 μM) at the indicated times. Wells that did not respond to injection and edge wells were omitted from analysis.

### Flow Cytometry

On a Matrigel-coated six-well dishes, 600,000 cells/well were plated and incubated for 3 d. Cells were infected with HCMV at an MOI of 0.05 IU/cell, washed at 2 hpi, and treated with 200 uM DETA/NO, DETA (spent control), or vehicle control. Treatment was changed every 2 d. At 11 dpi, cells were collected by trypsinization, washed with PBS, and fixed and permeabilized using Cell Fixation and Permeabilization Kit (Abcam) at room temp. Cells were incubated with conjugated antibody for 30 min at room temp in the dark and strained through 100 or 40 μM filter. Data was acquired on a BD LSR II flow cytometer (BD Biosciences). Cells were gated as described in the figure legends.

For flow cytometry, the following antibodies were used: Mouse anti-Nestin conjugated to V450 (clone 25, 2.5 μl, BD Biosciences), mouse anti-GFAP conjugated to PE (clone 1B4; 2.5 μl, BD Biosciences), mouse anti-Beta Tubulin Class III (clone Tuj1, 15 μl, BD Biosciences).

### Immunofluorescence

For organoid images, cortical organoids were fixed in 4% paraformaldehyde overnight at 4°C, washed with PBS, placed in 30% sucrose in PBS for two-three days, and then transferred to optimum cutting temperature (OCT) compound. Organoids were embedded in a cryosectioning mold using OCT compound (ThermoFisher), froze on dry ice, and cryosectioned into 20 μm slices. Sections were immediately mounted and allowed to dry. Sections were blocked in 5% donkey serum and 0.1% Triton-X100 in PBS for 30 min, incubated in primary antibody overnight at 4°C, and incubated in secondary antibody for 1 h at room temp. Hoechst was used to stain nuclei. Images were acquired on an upright TS100 Nikon fluorescence microscope and NIS Elements was used for imaging and analysis.

For NPCs, 600,000 cells/well were plated on a Matrigel-coated six-well dish containing 12mm coverslips and incubated for 3 d. Cells were infected with HCMV at an MOI of 0.05 IU/cell, washed at 2 hpi, and treated with 200 uM DETA/NO, DETA (spent control), or vehicle control. Treatment was changed every 2 d. At 11 dpi, cells were fixed in 4% paraformaldehyde for 20 min and permeabilized in 0.1% Triton X-100 for 15 min. Cells were blocked in 5% normal goat serum (Millipore-Sigma) in 0.2% Triton X-100 for 1 hour and incubated in primary antibody diluted in 5% normal goat serum in 0.1% Triton X-100 overnight at 4°C. Cells were washed with PBS-T and incubated for 1 hour with appropriate secondary antibody conjugated to Alexa Fluor 568 (1:1000) diluted in 5% normal goat serum in 0.1% Triton X-100 at room temperature. Cells were stained with Hoechst (1:1000) for 10 min at room temperature in dark. Coverslips were placed on glass slides and mounted with ProLong Glass Antifade Mountant. Images were acquired on a Nikon Ti Eclipse inverted microscope and NIS Elements was used for imaging and analysis.

The following antibodies were used for immunofluorescence: Mouse anti-Nestin (1:600; Abcam), rabbit anti-GFAP (1:1000; Agilent), mouse anti-Tuj1(1:100-1000; Millipore-Sigma), rabbit anti-SOX2 (Millipore Sigma; 1:100), anti-mouse Alexa Fluor-568 (1:1000; ThermoFisher), anti-rabbit Alexa Fluor-568 (1:1000; ThermoFisher), anti-mouse Alexa Fluor-647 (1:250), and Hoechst (1:1000; ThermoFisher). TUNEL assay was performed according to manufacturer’s instructions (ThermoFisher).

### Calcium imaging

On a Matrigel-coated six-well dish containing 12mm coverslips, 600,000 cells/well were plated and incubated for 3 d. Cells were infected with HCMV at an MOI of 0.05 IU/cell or mock-infected, washed at 2 hpi, and treated with 200 uM DETA/NO, DETA (spent control), or vehicle control. Treatment was changed every 2 d. At 11 dpi, cells were subject to Ca^2+^ imaging as described in Barabas et al. (94) and McGivern et al. (95). Briefly, live-cell calcium imaging was performed using the ratiometric dual-fluorescent calcium indicator FURA-2AM (ThermoFisher). Coverslips were loaded with 2.5 μl of FURA-2AM in 2% bovine serum albumin (BSA) in extracellular normal HEPES (ENH) buffer (150 μM NaCl, 10 μM HEPES, 8 μM glucose, 5.6 μM KCl, 2 μM CaCl2, 1 μM MgCl2) for 1 h, washed with ENH buffer for 20 min, and mounted onto a perfusion chamber. Brightfield and fluorescent images were taken prior to recordings. Coverslips were super fused with ENH buffer at 6 ml/min for 1 min prior to stimulation. To stimulate purinergic responses, coverslips were superfused with 10 μM ATP in ENH buffer for 1 min starting at 100 s after the start of recording. To stimulate voltage-gated channels, coverslips were super fused with 50 mM KCl in ENH buffer for 30 s at 175 s after the start of recording. Cells were washed with buffer between stimulations, with average baseline levels determined 30 s prior to each stimulation. NIS Elements (Nikon) was used for image acquisition and analysis. A cell selection tool was used to record calcium signals from individual cells, and 3 ROIs each containing 100 cells were analyzed per NPC coverslip. This method was repeated three times for each experimental group. Data are plotted as the ratio of bound (340 nm) to unbound (380 nm) intracellular Ca2^+^ over time in seconds.

### Statistical Analysis

Statistical significance was determined using Graphpad Prism application software and the appropriate statistical tests as indicated in the figure legends. *P* < 0.05 was considered significant.

## Acknowledgments

We thank Tom Shenk for providing antibodies against HCMV IE1 protein; Monika Zielonka in the Redox & Bioenergetics Shared Resource facility; Benedetta Bonacci in the Versiti-Blood Research Institute Flow Cytometry Core; and MCW Cancer Center and Children’s Research Institute’s Flow Cytometry Core. We thank Melissa Whyte and Halli Miller for their helpful advice on flow cytometry, Megan Schumacher for assistance with cell culture, Michele Battle for review of the manuscript, and members of the Terhune and Hudson laboratories for their input on the project.

Research reported in this publication was supported by the National Institute of Allergy and Infectious Diseases division of the National Institutes of Health under award number R01AI083281 (S.S.T.) and R01AI132414 (S.S.T. and A.D.E.). The content is solely the responsibility of the authors and does not necessarily represent the official views of the National Institutes of Health.

Author contributions were as follows: R.L.M., A.D.E., and S.S.T., conceptualization and methodology; S.S.T., A.D.E., funding acquisition and supervision; R.L.M., B.S.O., J.W.A., S.R., investigation; R.L.M., B.S.O., J.W.A., S.R., A.D.E., S.S.T., formal analysis; R.L.M. and S.S.T., visualization; R.L.M., writing—original draft; R.L.M., A.D.E., and S.S.T., writing—review & editing.

---

**Fig S1.**
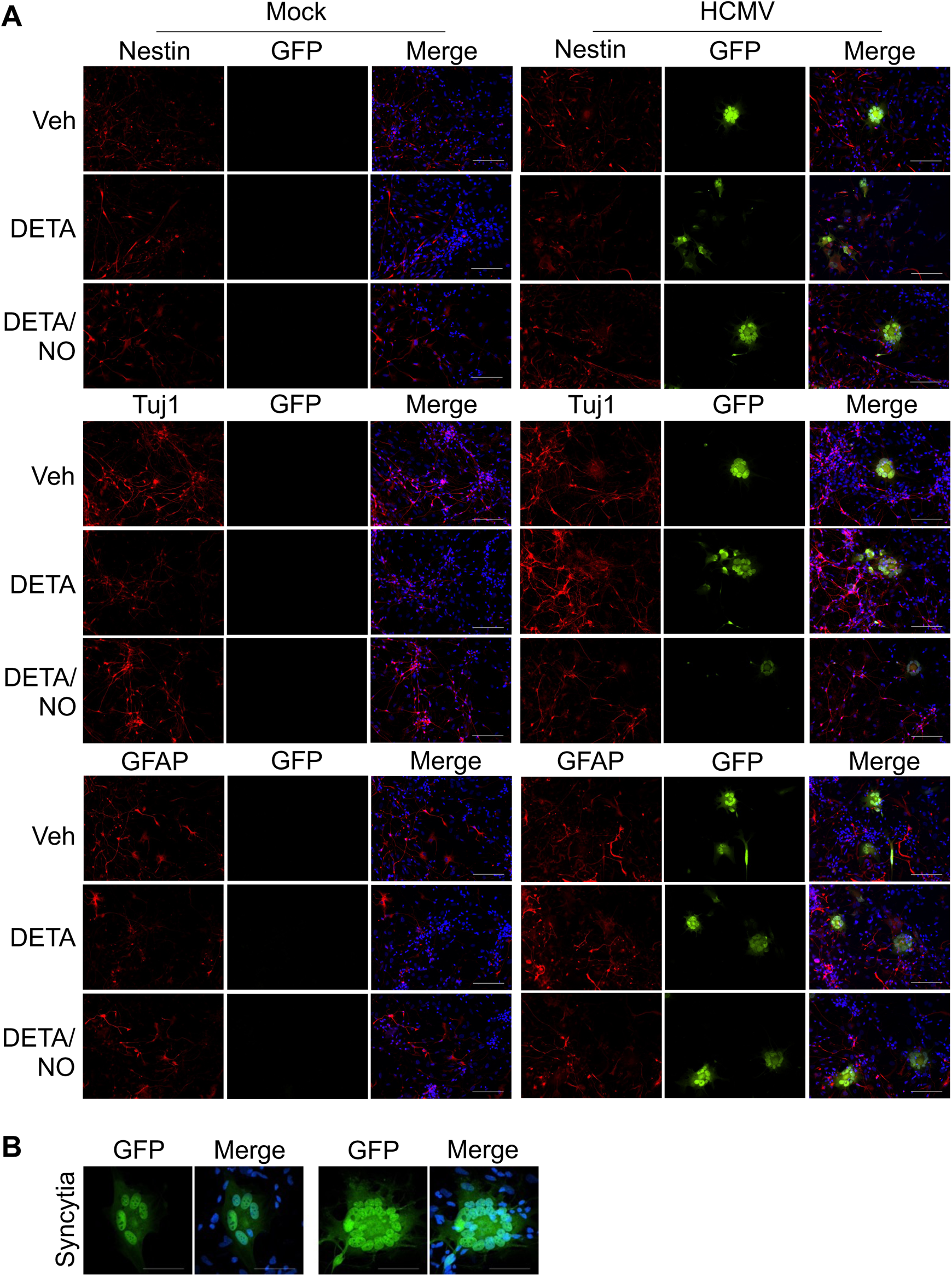
Assessing NPC differentiation using immunofluorescence. **(A)** Following the experimental timeline defined in Fig 6B, progenitor, glial, and neuron populations were assessed using immunofluorescence and labeling for Nestin, GFAP, and Tuj1, respectively. Data represent two biological replicates. Scale bar is 100 μm. **(B)** Multi-nucleated aggregate or syncytia formation in differentiated neural cells at 11 dpi. Images are maximum intensity projection. Scale bar is 50 μM.

**Fig S2.**
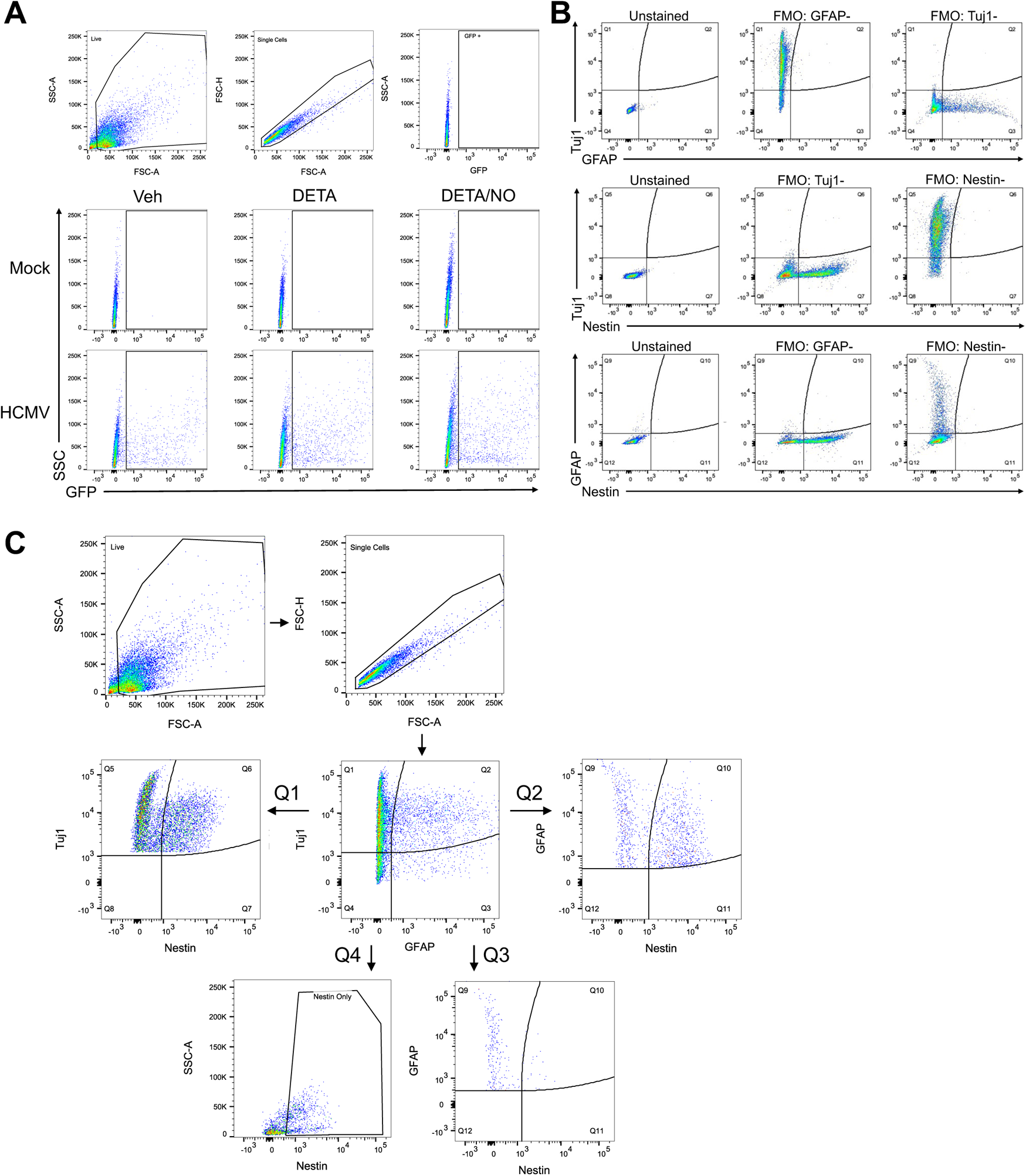
Gating for GFP and flow controls. **(A)** Live, single cell events were gated and the GFP-positive (GFP+) placed on the negative population of uninfected, unstained cells. This gate was applied to both mock and HCMV-infected samples for all conditions. **(B)** Fluorescence minus one (FMO) controls for flow analysis using unstained cells and FMO controls labeled with two of the three antibodies: Tuj1 and Nestin, Tuj1 and GFAP, and Nestin and GFAP. Live, single cell events were gated and FMO gates were set using the negative populations for each marker. **(C)** Live, single cells were gated and Tuj1+Nestin+GFAP- and Tuj1+Nestin-GFAP-events were quantified from Tuj1+ events gated from Tuj1/GFAP populations (Q1). Tuj1+Nestin-GFAP+ and Tuj1+Nestin+GFAP+ populations were quantified from cells gated on Tuj1+GFAP+ events (Q2). Tuj1-Nestin-GFAP+ populations were quantified by gating on Tuj1-GFAP+ events (Q3). Tuj1-Nestin+GFAP-populations were measured from gating on Tuj1-GFAP-events (Q4) and then Nestin+ events.

## References

1. Shenk TE, Stinski MF. 2008. Human cytomegalovirus. Preface. Curr Top Microbiol Immunol 325:v.

2. Pass RF, Arav-Boger R. 2018. Maternal and fetal cytomegalovirus infection: diagnosis, management, and prevention. F1000Res 7:255.

3. Lombardi G, Garofoli F, Stronati M. 2010. Congenital cytomegalovirus infection: treatment, sequelae and follow-up. J Matern Fetal Neonatal Med 23 Suppl 3:45–8.

4. Boppana SB, Ross SA, Fowler KB. 2013. Congenital cytomegalovirus infection: clinical outcome. Clin Infect Dis 57 Suppl 4:S178–81.

5. Dollard SC, Grosse SD, Ross DS. 2007. New estimates of the prevalence of neurological and sensory sequelae and mortality associated with congenital cytomegalovirus infection. Rev Med Virol 17:355–63.

6. Gabrielli L, Bonasoni MP, Lazzarotto T, Lega S, Santini D, Foschini MP, Guerra B, Baccolini F, Piccirilli G, Chiereghin A, Petrisli E, Gardini G, Lanari M, Landini MP. 2009. Histological findings in foetuses congenitally infected by cytomegalovirus. J Clin Virol 46 Suppl 4:S16–21.

7. Gabrielli L, Bonasoni MP, Santini D, Piccirilli G, Chiereghin A, Petrisli E, Dolcetti R, Guerra B, Piccioli M, Lanari M, Landini MP, Lazzarotto T. 2012. Congenital cytomegalovirus infection: patterns of fetal brain damage. Clin Microbiol Infect 18:E419–27.

8. Teissier N, Fallet-Bianco C, Delezoide AL, Laquerriere A, Marcorelles P, Khung-Savatovsky S, Nardelli J, Cipriani S, Csaba Z, Picone O, Golden JA, Van Den Abbeele T, Gressens P, Adle-Biassette H. 2014. Cytomegalovirus-induced brain malformations in fetuses. J Neuropathol Exp Neurol 73:143–58.

9. Sinzger C, Jahn G. 1996. Human cytomegalovirus cell tropism and pathogenesis. Intervirology 39:302–19.

10. Martinez-Cerdeno V, Noctor SC. 2018. Neural Progenitor Cell Terminology. Front Neuroanat 12:104.

11. Pan X, Li XJ, Liu XJ, Yuan H, Li JF, Duan YL, Ye HQ, Fu YR, Qiao GH, Wu CC, Yang B, Tian XH, Hu KH, Miao LF, Chen XL, Zheng J, Rayner S, Schwartz PH, Britt WJ, Xu J, Luo MH. 2013. Later passages of neural progenitor cells from neonatal brain are more permissive for human cytomegalovirus infection. J Virol 87:10968–79.

12. Gonzalez-Sanchez HM, Monsivais-Urenda A, Salazar-Aldrete CA, Hernandez-Salinas A, Noyola DE, Jimenez-Capdeville ME, Martinez-Serrano A, Castillo CG. 2015. Effects of cytomegalovirus infection in human neural precursor cells depend on their differentiation state. J Neurovirol 21:346–57.

13. Luo MH, Hannemann H, Kulkarni AS, Schwartz PH, O’Dowd JM, Fortunato EA. 2010. Human cytomegalovirus infection causes premature and abnormal differentiation of human neural progenitor cells. J Virol 84:3528–41.

14. Liu XJ, Yang B, Huang SN, Wu CC, Li XJ, Cheng S, Jiang X, Hu F, Ming YZ, Nevels M, Britt WJ, Rayner S, Tang Q, Zeng WB, Zhao F, Luo MH. 2017. Human cytomegalovirus IE1 downregulates Hes1 in neural progenitor cells as a potential E3 ubiquitin ligase. PLoS Pathog 13:e1006542.

15. Odeberg J, Wolmer N, Falci S, Westgren M, Seiger A, Soderberg-Naucler C. 2006. Human cytomegalovirus inhibits neuronal differentiation and induces apoptosis in human neural precursor cells. J Virol 80:8929–39.

16. Bigley TM, McGivern JV, Ebert AD, Terhune SS. 2016. Impact of a cytomegalovirus kinase inhibitor on infection and neuronal progenitor cell differentiation. Antiviral Res 129:67–73.

17. Reitsma JM, Sato H, Nevels M, Terhune SS, Paulus C. 2013. Human cytomegalovirus IE1 protein disrupts interleukin-6 signaling by sequestering STAT3 in the nucleus. J Virol 87:10763–76.

18. Harwardt T, Lukas S, Zenger M, Reitberger T, Danzer D, Ubner T, Munday DC, Nevels M, Paulus C. 2016. Human Cytomegalovirus Immediate-Early 1 Protein Rewires Upstream STAT3 to Downstream STAT1 Signaling Switching an IL6-Type to an IFNgamma-Like Response. PLoS Pathog 12:e1005748.

19. Wu CC, Jiang X, Wang XZ, Liu XJ, Li XJ, Yang B, Ye HQ, Harwardt T, Jiang M, Xia HM, Wang W, Britt WJ, Paulus C, Nevels M, Luo MH. 2018. Human Cytomegalovirus Immediate Early 1 Protein Causes Loss of SOX2 from Neural Progenitor Cells by Trapping Unphosphorylated STAT3 in the Nucleus. J Virol 92.

20. Sison SL, O’Brien BS, Johnson AJ, Seminary ER, Terhune SS, Ebert AD. 2019. Human Cytomegalovirus Disruption of Calcium Signaling in Neural Progenitor Cells and Organoids. J Virol 93.

21. Chiaradia I, Lancaster MA. 2020. Brain organoids for the study of human neurobiology at the interface of in vitro and in vivo. Nat Neurosci 23:1496–1508.

22. Brown RM, Rana P, Jaeger HK, O’Dowd JM, Balemba OB, Fortunato EA. 2019. Human Cytomegalovirus Compromises Development of Cerebral Organoids. J Virol 93.

23. Sun G, Chiuppesi F, Chen X, Wang C, Tian E, Nguyen J, Kha M, Trinh D, Zhang H, Marchetto MC, Song H, Ming GL, Gage FH, Diamond DJ, Wussow F, Shi Y. 2020. Modeling Human Cytomegalovirus-Induced Microcephaly in Human iPSC-Derived Brain Organoids. Cell Rep Med 1:100002.

24. MacMicking J, Xie QW, Nathan C. 1997. Nitric oxide and macrophage function. Annu Rev Immunol 15:323–50.

25. Boje KM, Arora PK. 1992. Microglial-produced nitric oxide and reactive nitrogen oxides mediate neuronal cell death. Brain Res 587:250–6.

26. Hall CN, Garthwaite J. 2009. What is the real physiological NO concentration in vivo? Nitric Oxide 21:92–103.

27. Jimenez W, Ros J, Morales-Ruiz M, Navasa M, Sole M, Colmenero J, Sort P, Rivera F, Arroyo V, Rodes J. 1999. Nitric oxide production and inducible nitric oxide synthase expression in peritoneal macrophages of cirrhotic patients. Hepatology 30:670–6.

28. Poderoso JJ, Carreras MC, Lisdero C, Riobo N, Schopfer F, Boveris A. 1996. Nitric oxide inhibits electron transfer and increases superoxide radical production in rat heart mitochondria and submitochondrial particles. Arch Biochem Biophys 328:85–92.

29. Drapier JC, Hibbs JB, Jr. 1986. Murine cytotoxic activated macrophages inhibit aconitase in tumor cells. Inhibition involves the iron-sulfur prosthetic group and is reversible. J Clin Invest 78:790–7.

30. Iglesias DE, Bombicino SS, Valdez LB, Boveris A. 2015. Nitric oxide interacts with mitochondrial complex III producing antimycin-like effects. Free Radic Biol Med 89:602–13.

31. Carreira BP, Morte MI, Inacio A, Costa G, Rosmaninho-Salgado J, Agasse F, Carmo A, Couceiro P, Brundin P, Ambrosio AF, Carvalho CM, Araujo IM. 2010. Nitric oxide stimulates the proliferation of neural stem cells bypassing the epidermal growth factor receptor. Stem Cells 28:1219–30.

32. Carreira BP, Morte MI, Santos AI, Lourenco AS, Ambrosio AF, Carvalho CM, Araujo IM. 2014. Nitric oxide from inflammatory origin impairs neural stem cell proliferation by inhibiting epidermal growth factor receptor signaling. Front Cell Neurosci 8:343.

33. Bergsland M, Covacu R, Perez Estrada C, Svensson M, Brundin L. 2014. Nitric oxide-induced neuronal to glial lineage fate-change depends on NRSF/REST function in neural progenitor cells. Stem Cells 32:2539–49.

34. Reiss CS, Komatsu T. 1998. Does nitric oxide play a critical role in viral infections? J Virol 72:4547–51.

35. Croen KD. 1993. Evidence for antiviral effect of nitric oxide. Inhibition of herpes simplex virus type 1 replication. J Clin Invest 91:2446–52.

36. Benencia F, Gamba G, Cavalieri H, Courreges MC, Benedetti R, Villamil SM, Massouh EJ. 2003. Nitric oxide and HSV vaginal infection in BALB/c mice. Virology 309:75–84.

37. Mannick JB, Asano K, Izumi K, Kieff E, Stamler JS. 1994. Nitric oxide produced by human B lymphocytes inhibits apoptosis and Epstein-Barr virus reactivation. Cell 79:1137–46.

38. Gao X, Tajima M, Sairenji T. 1999. Nitric oxide down-regulates Epstein-Barr virus reactivation in epithelial cell lines. Virology 258:375–81.

39. Cymerys J, Kowalczyk A, Mikolajewicz K, Slonska A, Krzyzowska M. 2019. Nitric Oxide Influences HSV-1-Induced Neuroinflammation. Oxid Med Cell Longev 2019:2302835.

40. Getts DR, Terry RL, Getts MT, Muller M, Rana S, Deffrasnes C, Ashhurst TM, Radford J, Hofer M, Thomas S, Campbell IL, King NJ. 2012. Targeted blockade in lethal West Nile virus encephalitis indicates a crucial role for very late antigen (VLA)-4-dependent recruitment of nitric oxide-producing macrophages. J Neuroinflammation 9:246.

41. Wang J, Liu J, Zhou R, Ding X, Zhang Q, Zhang C, Li L. 2018. Zika virus infected primary microglia impairs NPCs proliferation and differentiation. Biochem Biophys Res Commun 497:619–625.

42. Noda S, Tanaka K, Sawamura S, Sasaki M, Matsumoto T, Mikami K, Aiba Y, Hasegawa H, Kawabe N, Koga Y. 2001. Role of nitric oxide synthase type 2 in acute infection with murine cytomegalovirus. J Immunol 166:3533–41.

43. Dighiero P, Reux I, Hauw JJ, Fillet AM, Courtois Y, Goureau O. 1994. Expression of inducible nitric oxide synthase in cytomegalovirus-infected glial cells of retinas from AIDS patients. Neurosci Lett 166:31–4.

44. Hsu WM, Chen SS, Peng CH, Chen CF, Ko YC, Tsai DC, Chou CK, Ho LL, Chiou SH, Liu JH. 2003. Elevated nitric oxide level in aqueous humor of AIDS patients with cytomegalovirus retinitis. Ophthalmologica 217:298–301.

45. Yaiw KC, Mohammad AA, Taher C, Wilhelmi V, Davoudi B, Straat K, Assinger A, Ovchinnikova O, Shlyakhto E, Rahbar A, Koutonguk O, Religa P, Butler L, Khan Z, Streblow D, Pernow J, Soderberg-Naucler C. 2014. Human cytomegalovirus induces upregulation of arginase II: possible implications for vasculopathies. Basic Res Cardiol 109:401.

46. Nukui M, Roche KL, Jia J, Fox PL, Murphy EA. 2020. Protein S-Nitrosylation of Human Cytomegalovirus pp71 Inhibits Its Ability To Limit STING Antiviral Responses. J Virol 94.

47. Drutman SB, Mansouri D, Mahdaviani SA, Neehus AL, Hum D, Bryk R, Hernandez N, Belkaya S, Rapaport F, Bigio B, Fisch R, Rahman M, Khan T, Al Ali F, Marjani M, Mansouri N, Lorenzo-Diaz L, Emile JF, Marr N, Jouanguy E, Bustamante J, Abel L, Boisson-Dupuis S, Beziat V, Nathan C, Casanova JL. 2020. Fatal Cytomegalovirus Infection in an Adult with Inherited NOS2 Deficiency. N Engl J Med 382:437–445.

48. Mokry RL, Schumacher ML, Hogg N, Terhune SS. 2020. Nitric Oxide Circumvents Virus-Mediated Metabolic Regulation during Human Cytomegalovirus Infection. mBio 11.

49. Sakao-Suzuki M, Kawasaki H, Akamatsu T, Meguro S, Miyajima H, Iwashita T, Tsutsui Y, Inoue N, Kosugi I. 2014. Aberrant fetal macrophage/microglial reactions to cytomegalovirus infection. Ann Clin Transl Neurol 1:570–88.

50. Kosugi I, Kawasaki H, Arai Y, Tsutsui Y. 2002. Innate immune responses to cytomegalovirus infection in the developing mouse brain and their evasion by virus-infected neurons. Am J Pathol 161:919–28.

51. Keefer LK, Nims RW, Davies KM, Wink DA. 1996. “NONOates” (1-substituted diazen-1-ium-1,2-diolates) as nitric oxide donors: convenient nitric oxide dosage forms. Methods Enzymol 268:281–93.

52. Ebert AD, Yu J, Rose FF, Jr., Mattis VB, Lorson CL, Thomson JA, Svendsen CN. 2009. Induced pluripotent stem cells from a spinal muscular atrophy patient. Nature 457:277–80.

53. Consortium HDi. 2012. Induced pluripotent stem cells from patients with Huntington’s disease show CAG-repeat-expansion-associated phenotypes. Cell Stem Cell 11:264–78.

54. Lancaster MA, Renner M, Martin CA, Wenzel D, Bicknell LS, Hurles ME, Homfray T, Penninger JM, Jackson AP, Knoblich JA. 2013. Cerebral organoids model human brain development and microcephaly. Nature 501:373–9.

55. Zhou C, Yang X, Sun Y, Yu H, Zhang Y, Jin Y. 2016. Comprehensive profiling reveals mechanisms of SOX2-mediated cell fate specification in human ESCs and NPCs. Cell Res 26:171–89.

56. Miyagi S, Masui S, Niwa H, Saito T, Shimazaki T, Okano H, Nishimoto M, Muramatsu M, Iwama A, Okuda A. 2008. Consequence of the loss of Sox2 in the developing brain of the mouse. FEBS Lett 582:2811–5.

57. Favaro R, Valotta M, Ferri AL, Latorre E, Mariani J, Giachino C, Lancini C, Tosetti V, Ottolenghi S, Taylor V, Nicolis SK. 2009. Hippocampal development and neural stem cell maintenance require Sox2-dependent regulation of Shh. Nat Neurosci 12:1248–56.

58. Graham V, Khudyakov J, Ellis P, Pevny L. 2003. SOX2 functions to maintain neural progenitor identity. Neuron 39:749–65.

59. Ebert AD, Shelley BC, Hurley AM, Onorati M, Castiglioni V, Patitucci TN, Svendsen SP, Mattis VB, McGivern JV, Schwab AJ, Sareen D, Kim HW, Cattaneo E, Svendsen CN. 2013. EZ spheres: a stable and expandable culture system for the generation of pre-rosette multipotent stem cells from human ESCs and iPSCs. Stem Cell Res 10:417–427.

60. Wang D, Yu QC, Schroer J, Murphy E, Shenk T. 2007. Human cytomegalovirus uses two distinct pathways to enter retinal pigmented epithelial cells. Proc Natl Acad Sci U S A 104:20037–42.

61. Takahashi J, Palmer TD, Gage FH. 1999. Retinoic acid and neurotrophins collaborate to regulate neurogenesis in adult-derived neural stem cell cultures. J Neurobiol 38:65–81.

62. Seoane J, Le HV, Shen L, Anderson SA, Massague J. 2004. Integration of Smad and forkhead pathways in the control of neuroepithelial and glioblastoma cell proliferation. Cell 117:211–23.

63. Lodato MA, Ng CW, Wamstad JA, Cheng AW, Thai KK, Fraenkel E, Jaenisch R, Boyer LA. 2013. SOX2 co-occupies distal enhancer elements with distinct POU factors in ESCs and NPCs to specify cell state. PLoS Genet 9:e1003288.

64. Maffezzini C, Calvo-Garrido J, Wredenberg A, Freyer C. 2020. Metabolic regulation of neurodifferentiation in the adult brain. Cell Mol Life Sci 77:2483–2496.

65. Zheng X, Boyer L, Jin M, Mertens J, Kim Y, Ma L, Ma L, Hamm M, Gage FH, Hunter T. 2016. Metabolic reprogramming during neuronal differentiation from aerobic glycolysis to neuronal oxidative phosphorylation. Elife 5.

66. Osborne GW. 2013. Flow cytometry of neural cells. Methods Mol Biol 1059:135–44.

67. Rieske P, Azizi SA, Augelli B, Gaughan J, Krynska B. 2007. A population of human brain parenchymal cells express markers of glial, neuronal and early neural cells and differentiate into cells of neuronal and glial lineages. Eur J Neurosci 25:31–7.

68. Draberova E, Del Valle L, Gordon J, Markova V, Smejkalova B, Bertrand L, de Chadarevian JP, Agamanolis DP, Legido A, Khalili K, Draber P, Katsetos CD. 2008. Class III beta-tubulin is constitutively coexpressed with glial fibrillary acidic protein and nestin in midgestational human fetal astrocytes: implications for phenotypic identity. J Neuropathol Exp Neurol 67:341–54.

69. Lin JH, Takano T, Arcuino G, Wang X, Hu F, Darzynkiewicz Z, Nunes M, Goldman SA, Nedergaard M. 2007. Purinergic signaling regulates neural progenitor cell expansion and neurogenesis. Dev Biol 302:356–66.

70. Spitzer NC, Root CM, Borodinsky LN. 2004. Orchestrating neuronal differentiation: patterns of Ca2+ spikes specify transmitter choice. Trends Neurosci 27:415–21.

71. Gu X, Spitzer NC. 1995. Distinct aspects of neuronal differentiation encoded by frequency of spontaneous Ca2+ transients. Nature 375:784–7.

72. Ormel PR, Vieira de Sa R, van Bodegraven EJ, Karst H, Harschnitz O, Sneeboer MAM, Johansen LE, van Dijk RE, Scheefhals N, Berdenis van Berlekom A, Ribes Martinez E, Kling S, MacGillavry HD, van den Berg LH, Kahn RS, Hol EM, de Witte LD, Pasterkamp RJ. 2018. Microglia innately develop within cerebral organoids. Nat Commun 9:4167.

73. Agostini M, Romeo F, Inoue S, Niklison-Chirou MV, Elia AJ, Dinsdale D, Morone N, Knight RA, Mak TW, Melino G. 2016. Metabolic reprogramming during neuronal differentiation. Cell Death Differ 23:1502–14.

74. Bifari F, Dolci S, Bottani E, Pino A, Di Chio M, Zorzin S, Ragni M, Zamfir RG, Brunetti D, Bardelli D, Delfino P, Cattaneo MG, Bordo R, Tedesco L, Rossi F, Bossolasco P, Corbo V, Fumagalli G, Nisoli E, Valerio A, Decimo I. 2020. Complete neural stem cell (NSC) neuronal differentiation requires a branched chain amino acids-induced persistent metabolic shift towards energy metabolism. Pharmacol Res 158:104863.

75. Li D, Ding Z, Gui M, Hou Y, Xie K. 2020. Metabolic Enhancement of Glycolysis and Mitochondrial Respiration Are Essential for Neuronal Differentiation. Cell Reprogram 22:291–299.

76. Brown GC. 2010. Nitric oxide and neuronal death. Nitric Oxide 23:153–65.

77. Toth AB, Shum AK, Prakriya M. 2016. Regulation of neurogenesis by calcium signaling. Cell Calcium 59:124–34.

78. Rosenberg SS, Spitzer NC. 2011. Calcium signaling in neuronal development. Cold Spring Harb Perspect Biol 3:a004259.

79. Sharon-Friling R, Goodhouse J, Colberg-Poley AM, Shenk T. 2006. Human cytomegalovirus pUL37x1 induces the release of endoplasmic reticulum calcium stores. Proc Natl Acad Sci U S A 103:19117–22.

80. McCormick AL, Smith VL, Chow D, Mocarski ES. 2003. Disruption of mitochondrial networks by the human cytomegalovirus UL37 gene product viral mitochondrion-localized inhibitor of apoptosis. J Virol 77:631–41.

81. Luganini A, Di Nardo G, Munaron L, Gilardi G, Fiorio Pla A, Gribaudo G. 2018. Human cytomegalovirus US21 protein is a viroporin that modulates calcium homeostasis and protects cells against apoptosis. Proc Natl Acad Sci U S A 115:E12370–E12377.

82. Wang X, Huong SM, Chiu ML, Raab-Traub N, Huang ES. 2003. Epidermal growth factor receptor is a cellular receptor for human cytomegalovirus. Nature 424:456–61.

83. Kaarbo M, Ager-Wick E, Osenbroch PO, Kilander A, Skinnes R, Muller F, Eide L. 2011. Human cytomegalovirus infection increases mitochondrial biogenesis. Mitochondrion 11:935–45.

84. Combs JA, Norton EB, Saifudeen ZR, Bentrup KHZ, Katakam PV, Morris CA, Myers L, Kaur A, Sullivan DE, Zwezdaryk KJ. 2020. Human Cytomegalovirus Alters Host Cell Mitochondrial Function during Acute Infection. J Virol 94.

85. Brown GC, Cooper CE. 1994. Nanomolar concentrations of nitric oxide reversibly inhibit synaptosomal respiration by competing with oxygen at cytochrome oxidase. FEBS Lett 356:295–8.

86. Chambers JW, Maguire TG, Alwine JC. 2010. Glutamine metabolism is essential for human cytomegalovirus infection. J Virol 84:1867–73.

87. Munger J, Bajad SU, Coller HA, Shenk T, Rabinowitz JD. 2006. Dynamics of the cellular metabolome during human cytomegalovirus infection. PLoS Pathog 2:e132.

88. Munger J, Bennett BD, Parikh A, Feng XJ, McArdle J, Rabitz HA, Shenk T, Rabinowitz JD. 2008. Systems-level metabolic flux profiling identifies fatty acid synthesis as a target for antiviral therapy. Nat Biotechnol 26:1179–86.

89. Rodriguez-Sanchez I, Schafer XL, Monaghan M, Munger J. 2019. The Human Cytomegalovirus UL38 protein drives mTOR-independent metabolic flux reprogramming by inhibiting TSC2. PLoS Pathog 15:e1007569.

90. Baldick CJ, Jr., Marchini A, Patterson CE, Shenk T. 1997. Human cytomegalovirus tegument protein pp71 (ppUL82) enhances the infectivity of viral DNA and accelerates the infectious cycle. J Virol 71:4400–8.

91. Sinzger C, Hahn G, Digel M, Katona R, Sampaio KL, Messerle M, Hengel H, Koszinowski U, Brune W, Adler B. 2008. Cloning and sequencing of a highly productive, endotheliotropic virus strain derived from human cytomegalovirus TB40/E. J Gen Virol 89:359–368.

92. Umashankar M, Petrucelli A, Cicchini L, Caposio P, Kreklywich CN, Rak M, Bughio F, Goldman DC, Hamlin KL, Nelson JA, Fleming WH, Streblow DN, Goodrum F. 2011. A novel human cytomegalovirus locus modulates cell type-specific outcomes of infection. PLoS Pathog 7:e1002444.

93. Yu D, Smith GA, Enquist LW, Shenk T. 2002. Construction of a self-excisable bacterial artificial chromosome containing the human cytomegalovirus genome and mutagenesis of the diploid TRL/IRL13 gene. J Virol 76:2316–28.

94. Barabas ME, Kossyreva EA, Stucky CL. 2012. TRPA1 is functionally expressed primarily by IB4-binding, non-peptidergic mouse and rat sensory neurons. PLoS One 7:e47988.

95. McGivern JV, Patitucci TN, Nord JA, Barabas MA, Stucky CL, Ebert AD. 2013. Spinal muscular atrophy astrocytes exhibit abnormal calcium regulation and reduced growth factor production. Glia 61:1418–1428.

96. Hoops S, Sahle S, Gauges R, Lee C, Pahle J, Simus N, Singhal M, Xu L, Mendes P, Kummer U. 2006. COPASI--a COmplex PAthway SImulator. Bioinformatics 22:3067–74.

97. Lewis RS, Deen WM. 1994. Kinetics of the reaction of nitric oxide with oxygen in aqueous solutions. Chem Res Toxicol 7:568–74.

